# Bottom-up proteomics under acidic conditions using protease type XIII from Aspergillus saitoi

**DOI:** 10.1101/2025.06.01.657254

**Authors:** Ryota Tomioka, Ayana Tomioka, Kosuke Ogata, Yasushi Ishihama

**Affiliations:** Graduate School of Pharmaceutical Sciences, Kyoto University, Kyoto 606-8501, Japan; Shionogi & Co., Ltd., Biopharmaceutical Research Division, Toyonaka, Osaka 561-0825, Japan; Laboratory of Proteomics for Drug Discovery, National Institute of Biomedical Innovation, Health and Nutrition, Ibaraki, Osaka, 567-0085, Japan

**Keywords:** Protease type XIII, Bottom-up proteomics, Protease, Deamidation, Cleavage preference

## Abstract

Bottom-up proteomics is a powerful technique for comprehensive analysis of proteins by proteolytic cleavage followed by liquid chromatography/tandem mass spectrometry to identify the resulting peptides. Trypsin is the gold-standard protease for bottom-up proteomics, though its cleavage specificity limits peptide identification, depending on the protein sequence. In addition, its optimal pH is weakly alkaline, which can cause modification artifacts such as deamidation. We hypothesized that these limitations might be overcome by using protease type XIII (P13ase) from *Aspergillus saitoi*, which is active at low pH. P13ase has been used for protein structural analysis by hydrogen-deuterium exchange mass spectrometry, but its cleavage preferences have not been clarified. Here, we show that P13ase primarily cleaves the C-terminal side of Lys, Arg, and Leu, and the optimal P13ase digestion conditions for bottom-up proteomics are pH 3.5, 37°C for 60 min. Under these conditions, sequence coverage of more than 90% was achieved for several proteins in HeLa cell extracts, which is unachievable with trypsin. In addition, P13ase digestion reduced artifacts such as deamidation products generated by cyclization reactions and subsequent hydrolysis. These results indicate that P13ase is a promising new tool for precision proteomics.

## INTRODUCTION

Protein digestion is an important technique for characterizing proteins. For example, the sequence of an antibody drug can be identified by mass spectrometric analysis of peptides generated by cleaving peptide bonds(1). Indeed, protein digestion is a crucial and essential step not only in such single protein analysis, but also in bottom-up proteomics using LC/MS/MS, and stringent control is required to ensure accurate identification and quantification of each protein(2, 3). Trypsin has been widely used as the gold standard protease in proteomics due to its cleavage specificity, high digestion efficiency, and ready availability(4, 5). Trypsin can specifically cleave the C-terminal side of lysine and arginine, but the peptides generated by trypsin are sometimes excessively long or short, making it difficult to identify the entire sequence of proteins of interest(4, 5). Therefore, alternative proteases with different specificities, such as Lys-C, Lys-N, Asp-N, Glu- C, Arg-C, chymotrypsin, and ProAlanase, have been adopted for bottom-up proteomics(6, 7), and combinations of these proteases have been reported to improve coverage compared to trypsin alone. In addition, modifying the sequence of proteases can enhance their functionality. For example, methylating the lysine residues of trypsin has been reported to inhibit self-digestion, and modifying the Arg-C sequence has been shown to improve the cleavage specificity(8, 9). Furthermore, subtilisin, proteinase K, and thermolysin, which have relatively low cleavage specificity, have been proposed as proteases for rapid digestion(10). In addition to these alternative proteases, chemical modifications have been employed to expand the cleavage sites of trypsin(11–13). However, there is still a need for new methods to improve coverage.

The challenges in trypsin digestion are not limited to the length of the generated peptides. Another issue is the modification artifacts generated due to the optimal pH of trypsin (pH 7 to 9). Specifically, under these conditions, artifacts are produced by cyclization reactions followed by hydrolysis(14, 15). These artifacts can cause false-positive identifications and distort quantitative analysis, thereby complicating data interpretation. Various attempts have been made to overcome this problem. Liu et al.(15) demonstrated that digestion using Glu-C protease at pH 4.5 in ammonium acetate effectively removed deamidation artifacts. Cao et al.(16) developed a new low-pH peptide mapping protocol combining Lys-C and modified trypsin to enable qualitative and quantitative characterization of succinimidyl intermediates and deamidation products. However, lowering the digestion pH shifts the protease out of its optimal pH range, reducing digestion efficiency. There are also heat-treated acidic archaeal proteases that function optimally at pH 2–4 and exhibit high digestion efficiency(17, 18), but their cleavage specificity is relatively low, and their artifact suppression efficacy remains unclear.

In trypsin digestion, the relatively long digestion time (overnight) is also an issue. While trypsin-immobilized spin columns can complete digestion in approximately 10 minutes(19–21), in-solution digestion is still superior in terms of cost and reproducibility, and methods to achieve short digestion times under in-solution digestion conditions are needed.

P13ase from *Aspergillus saitoi* digests a wide range of proteins with optimal activity at pH 2.5- 3.0(22, 23). It has received much attention in recent years, especially for application in HDX-MS, where it is essential to carry out digestion within a short time at low temperature and low pH(24–27). This protease is an acidic and non-specific protease, but its cleavage preference is not the same as that of pepsin, enabling the generation of a variety of peptides(26). While P13ase has been increasingly used in HDX-MS in recent years, it has hardly been utilized in bottom-up proteomics. One reason for this is that optimal digestion conditions, such as pH, temperature and digestion time, as well as the enzyme’s cleavage preferences, remain to be systematically evaluated. Understanding these parameters is critical to maximize the utility of type XIII proteases in a bottom-up proteomics workflow.

In this study, we comprehensively evaluated the proteolytic performance of P13ase using HeLa cell lysate as a model system, aiming to maximize the potential of P13ase with state-of-the-art LC/MS/MS analysis and data processing systems. The effects of digestion pH, temperature and reaction time on peptide and protein identification, the cleavage preferences and the extent of artifact formation were examined. Finally, P13ase was compared with trypsin under optimized conditions to establish its potential as a new protease for proteomics research.

## MATERIALS AND METHODS

### Materials

Urea, DTT, IAA, Ambic, ACN, acetic acid, sodium hydroxide, hydrochloric acid, formic acid and TFA were obtained from FUJIFILM Wako (Osaka, Japan). P13ase from *Aspergillus saitoi* was obtained from Sigma-Aldrich (St. Louis, MO). Sequencing-grade modified trypsin was obtained from Promega (Madison, WI). SDB-XC Empore disks were from GL Sciences (Tokyo, Japan). Water was purified with a PURELAB Quest system (ELGA LabWater, UK).

### Sample preparation

HeLa cells were cultured in Dulbecco’s modified Eagle’s medium containing 1× antibiotic-antimycotic solution with 10 % fetal bovine serum at 37 °C under 5 % CO2. HeLa cells were pelleted by centrifugation and proteins were denatured with 8 M urea in 50 mM Ambic solution and extracted by sonication. The concentration of proteins was determined by BCA assay. Proteins were reduced with 10 mM DTT in 50 mM Ambic solution at 37 °C for 30 min and carbamidomethylated with 50 mM IAA in 50 mM Ambic solution at 37 °C for 30 min. Proteins were 10-fold diluted with 1% formic acid (pH 1.5, 2.0, 2.5, 3.0, 3.5 and 4.0) for P13ase digestion or 10- fold diluted with 50 mM Ambic solution (pH 7.8) for trypsin digestion. Proteins were digested with P13ase (protein : enzyme = 10 : 1) or trypsin (protein : enzyme = 100 : 1). The enzyme-protein ratio for type XIII was established based on a previous report(28). Trypsin digestion was performed at 37°C for 16 hours, while the P13ase digestion conditions during the optimization process are described together with the results in the following section. Digests were desalted in reversed-phase StageTips(29, 30) as described previously.

### NanoLC/MS/MS

NanoLC/MS/MS was performed on an Ultimate 3000 liquid chromatograph combined with an Orbitrap Exploris 480 mass spectrometer (Thermo Fisher Scientific). The mobile phases for LC consisted of solvent A (0.5% acetic acid) and solvent B (0.5% acetic acid and 80% acetonitrile). Peptides were separated on self-pulled needle columns (250 mm length, 100 µm ID) packed with Reprosil-Pur 120 C18-AQ 1.9 µm reversed-phase material (Dr. Maisch, Ammerbuch, Germany) at 50 °C in a column oven (Sonation). The injection volume was 5 µL and the flow rate was 400 nL/min. Digested peptides were separated by applying a step gradient of 5 % B for 5 min, 5−19 % B for 55 min, 19−29 % B for 21 min, 29−40 % B for 9 min, 40-99 % B for 0.1 min and 99 % B for 5 min. Peptides were analyzed in the DDA mode. The electrospray voltage was set to 2.4 kV in the positive mode. The mass range of the survey scan was from 375 to 1500 *m/z* with a resolution of 60,000, 300% normalized AGC target, and auto maximum injection time. The first mass of MS/MS scan was set to 120 *m/z* with a resolution of 15,000, standard AGC target and auto maximum injection time. The fragmentation was performed by higher-energy collisional dissociation with a normalized collision energy of 30%. The dynamic exclusion time was set to 20 sec.

### Data Analysis and Bioinformatics

Peptides and proteins were identified through automated database searching using MSFragger(31, 32) version 4.1 and Philosopher(33) version 5.1 against the human database (release 2023/09) from UniProtKB/Swiss-Prot. Sequences of P13ase (UniProt Accession Q12567) and trypsin (UniProt Accession P00761) were used to evaluate self-digestion. For P13ase, the digestion mode was set to non-specific digestion. For trypsin, the digestion mode was set allowing for up to two missed cleavages. Oxidation (M), acetylation (protein N-term), deamidation (N) and NH3-loss (N) were allowed as variable modifications. Carbamidomethylation (C) was set as a fixed modification. The FDR filter was set to 0.01 at both the PSM and protein levels. Peptides and proteins were quantified by IonQuant(34). Searches were also performed in MaxQuant(35) version 2.4.2.0 for the analysis of cleavage preference. The cleavage preference was analyzed by iceLogo(36). The sequence coverages were calculated with Protein Coverage Summarizer (https://github.com/PNNL-Comp-Mass-Spec/protein-coverage-summarizer). Extracted ion chromatograms and mass spectra were analyzed by FreeStyle (Thermo Fisher Scientific). Motif analysis was performed using rmotif-X(37).

### Data availability

The MS raw data and analysis files have been deposited with the ProteomeXchange Consortium (https://proteomecentral.proteomexchange.org) via the jPOST partner repository (https:// jpostdb.org)(38) with the data set identifier PXD064095. Annotation of fragment ions for MS/MS spectra can be confirmed with PDV(39).

## RESULTS AND DISCUSSION

### Effect of pH on P13ase digestion

To evaluate the activity of P13ase in bottom-up proteomics, we first investigated the effect of pH. Based on previous reports indicating that the optimal pH for P13ase is 2.5–3.0 (23), we set the pH range for this study from 1.5 to 4.0. Proteins extracted from HeLa cells were used as samples and digested at 37 °C for 10 minutes. Triplicate experiments were conducted for each condition, and nanoLC/MS/MS measurements were performed in the DDA mode. Peptide identification was performed using MSFragger in the non-specific digestion mode. As a control, a sample was digested with trypsin under weakly alkaline conditions at 37 °C for 16 hours. The number of identified peptides and cleavage specificity are shown in Fig. 1 and Fig. 2, respectively. The number of identified peptides tended to increase with increasing pH. At pH 3.5, although the number of identified peptides was lower than in the case of trypsin digestion, 22,793 peptides and 2,334 proteins were identified, representing the highest number of unique peptides identified (Fig. 1A). The average length of identified peptides increased with decreasing pH at pH 2.5 or lower, but at pH 2.5 or higher, it remained at an average of approximately 14 amino acid residues. Compared with trypsin-digested peptides, they were approximately 2 amino acid residues longer (Fig. 1B). In reverse-phase LC separation, the retention time range showed a trend toward longer retention times at pH values of 2.5 or lower, but remained nearly constant at pH 2.5 or higher, differing from the peptide length pattern and exhibiting a retention time range similar to that of trypsin-digested peptides (Fig. 1C). The total ion current chromatograms showed that ions with longer retention times were more abundant at lower pH values, suggesting that digestion may not have been sufficient at lower pH (Supplemental Fig. S1).

**Figure 1:**
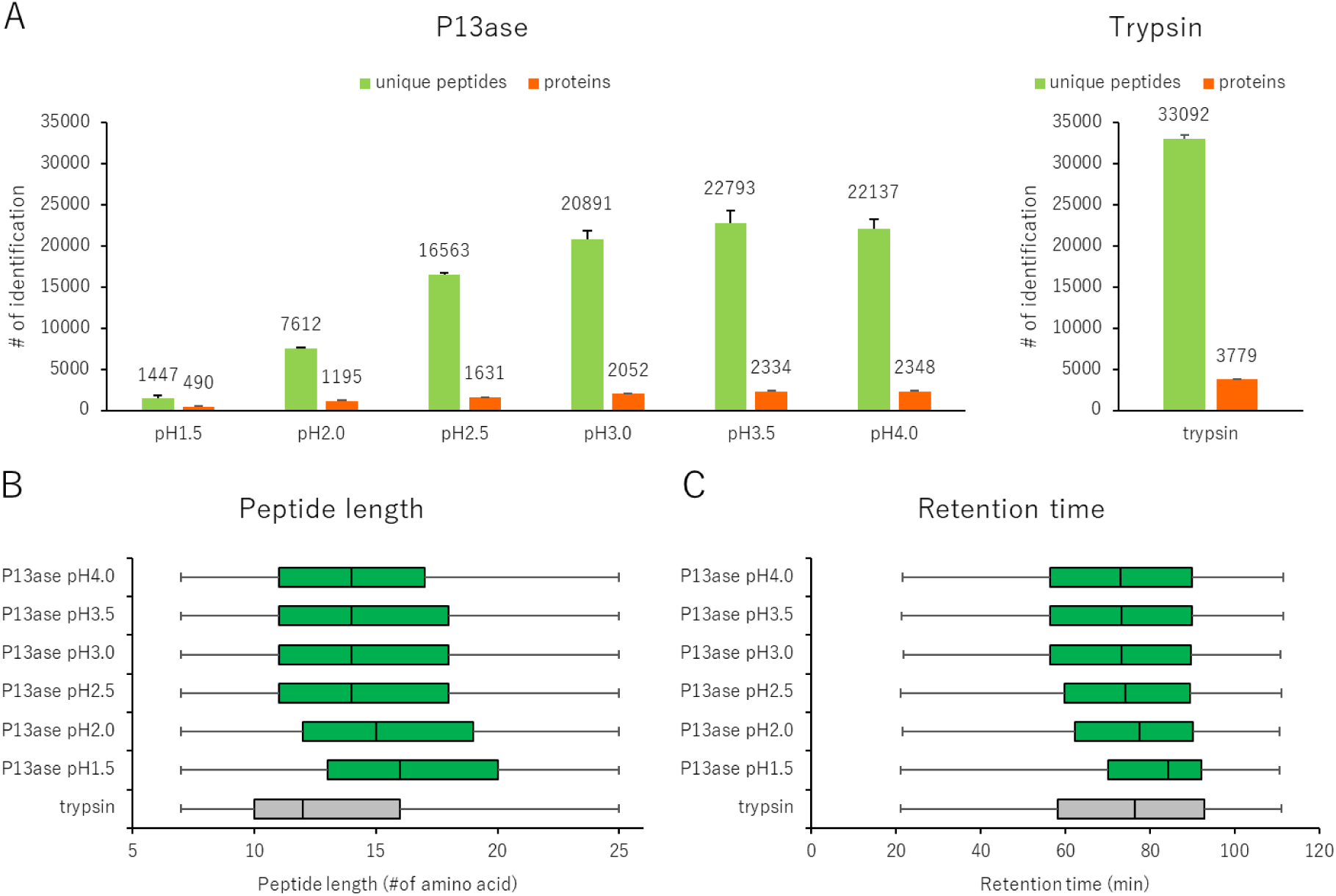
Effect of digestion pH on the activity of P13ase. (A) The number of unique peptides and proteins identified in HeLa cell extract at various pH values (1.5 ∼ 4.0). The bar graphs represent the mean of three replicates, and error bars represent the standard error. (B) The length distribution of identified peptides after P13ase digestion at various pH values. (C) The distribution of retention time of identified peptides after P13ase digestion at various pH values.

**Figure 2:**
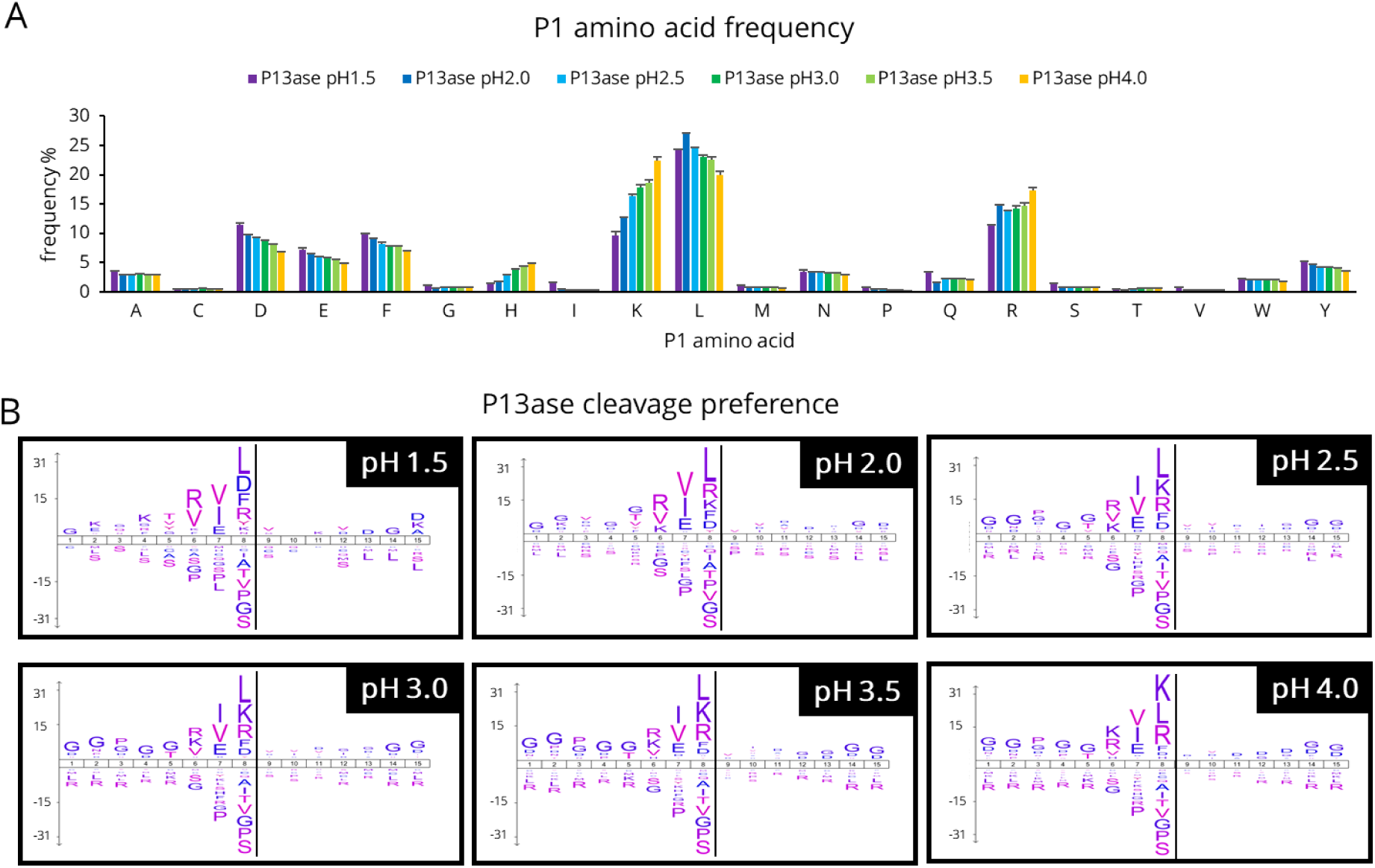
Cleavage specificity of P13ase in HeLa cell extract at various pH values. (A) Frequencies of P1 amino acids at cleavage sites at various pH values (1.5 ∼ 4.0). The bar graphs represent the mean of three replicates, and error bars represent the standard error. (B) Cleavage preferences visualized using iceLogo at various pH values (1.5 ∼ 4.0). Non-specific MaxQuant search was performed to calculate cleavage preferences against the HeLa cell lysate dataset (three replicates combined). The y-axis is the percentage difference in frequency for an amino acid in the experimental set and the reference set.

Using these identified peptides, the cleavage preferences were examined. Because previous reports on cleavage preferences of P13ase focused on the P1 position (26, 27), we first examined the amino acid frequency at the P1 position (Fig. 2A). As the pH was increased, the frequency of trypsin-like cleavage (Lys and Arg) increased, while the frequency of cleavage at acidic (Asp and Glu) and hydrophobic (Leu and Phe) amino acids decreased. Further analysis of the effect of pH on the frequency of missed cleavages showed that the percentage of peptides containing missed cleavages decreased at higher pH for Lys and Arg, whereas the decrease in missed cleavages was not significant for Leu, Phe, Asp, and Glu (Supplemental Fig. S2). Considering that the average peptide length at pH 2.5 and above was approximately 2 amino acid residues longer than tryptic peptides, even though there were other amino acids besides Lys and Arg that could be cleaved, this suggests that there is a contribution from sites other than the P1 position. Thus, we performed motif analysis of the cleavage preferences for 15 amino acid residues (from P8 to P7’) using iceLogo and motif-x (Fig. 2B and Supplemental Table S1). The results showed that not only P1 but also co-occurring sequences were observed, with a trend toward higher frequencies of Ile, Val, and Glu at P2 and Arg, Lys, and Val at P3. On the other hand, the reported hydrophobic amino acid preference at P1’(23) was not confirmed.

Compared to trypsin, P13ase is able to cleave the C-terminal side of Leu in addition to Lys and Arg, while cleavage at Ile does not occur at all. This result suggests that Leu, which cannot be distinguished from Ile by MS, may be unambiguously identified by P13ase digestion. Furthermore, it is known that under alkaline trypsin digestion conditions, Asn and Gln undergo cyclization reactions with the carbonyl groups of peptide bonds to produce succinimide intermediates followed by hydrolysis to produce deamidated peptides. Since the entire P13ase digestion process can be performed at low pH, one would expect these artificial modifications to be reduced. Indeed, the contents of both deamidated peptides and succinimide intermediates were significantly reduced by P13ase digestion compared to trypsin digestion under all pH conditions tested (Supplementary Fig. S3). This result will be discussed in more detail in the following section (Fig. 6D).

Based on these results, pH 3.5 was selected as the optimum pH for P13ase digestion in bottom-up proteomics.

### Effect of temperature and time on P13ase digestion

Next, we examined the digestion time (2 minutes to 16 hours) and temperature (4 °C, 25 °C, 37 °C) at pH 3.5. As expected, higher temperatures reduced the time required to maximize the number of identified peptides. At 37 °C for 60 minutes, 32,472 peptides and 2,721 proteins were identified, representing the highest number of unique peptides and proteins identified (Fig. 3A). Although the number of identified peptides and proteins was lower compared to trypsin, the number of peptides per identified protein for P13ase (11.6 peptides/protein) was higher than that for trypsin (9.8 peptides/protein). Additionally, the average length of the identified peptides showed a decreasing trend with increasing digestion time and temperature (Fig. 3B, Supplemental Fig. S4). Furthermore, the retention time range was similar to that of trypsin under most conditions, though in the case of 16 hours of digestion at 37 °C, the number of peptides with shorter retention times increased (Fig. 3C, Supplemental Fig. S5).

**Figure 3:**
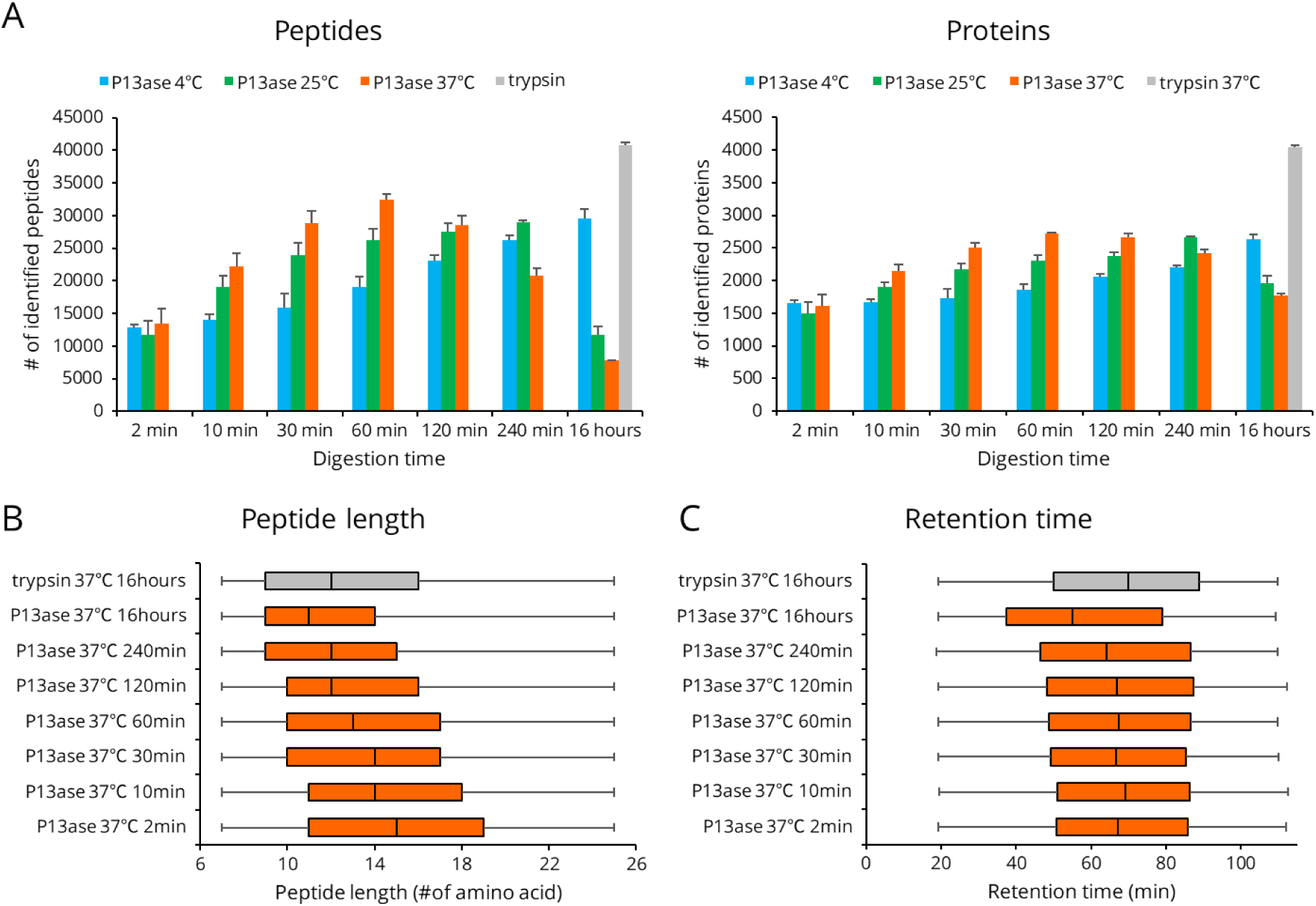
Effect of digestion temperature and time on the activity of P13ase in HeLa cell extract. (A) The number of identified unique peptides and proteins at various digestion temperature and times. The bar graphs represent the mean of three replicates, and error bars represent the standard error. (B) The length distribution of identified peptides after various digestion times at 37°C. (C) The distribution of retention times of identified peptides after various digestion times at 37°C.

When the cleavage preferences at 37 °C were examined, the cleavage preferences at positions P1, P2, and P3 were consistent with the results presented in the previous section for digestion at 60 min or less. However, the cleavage preferences at the P3 position disappeared when digestion was performed at 37 °C for 120 min or longer (Fig. 4A). A similar trend was observed when digestion was performed at 4 °C and 25 °C, and the lower the digestion temperature, the longer the digestion time required for the cleavage preferences to change (Supplemental Fig. S6). The number of motifs extracted by motif analysis was significantly reduced upon digestion for 16 hours, suggesting that more nonspecific cleavages occurred (Supplemental Table S2,3).

**Figure 4:**
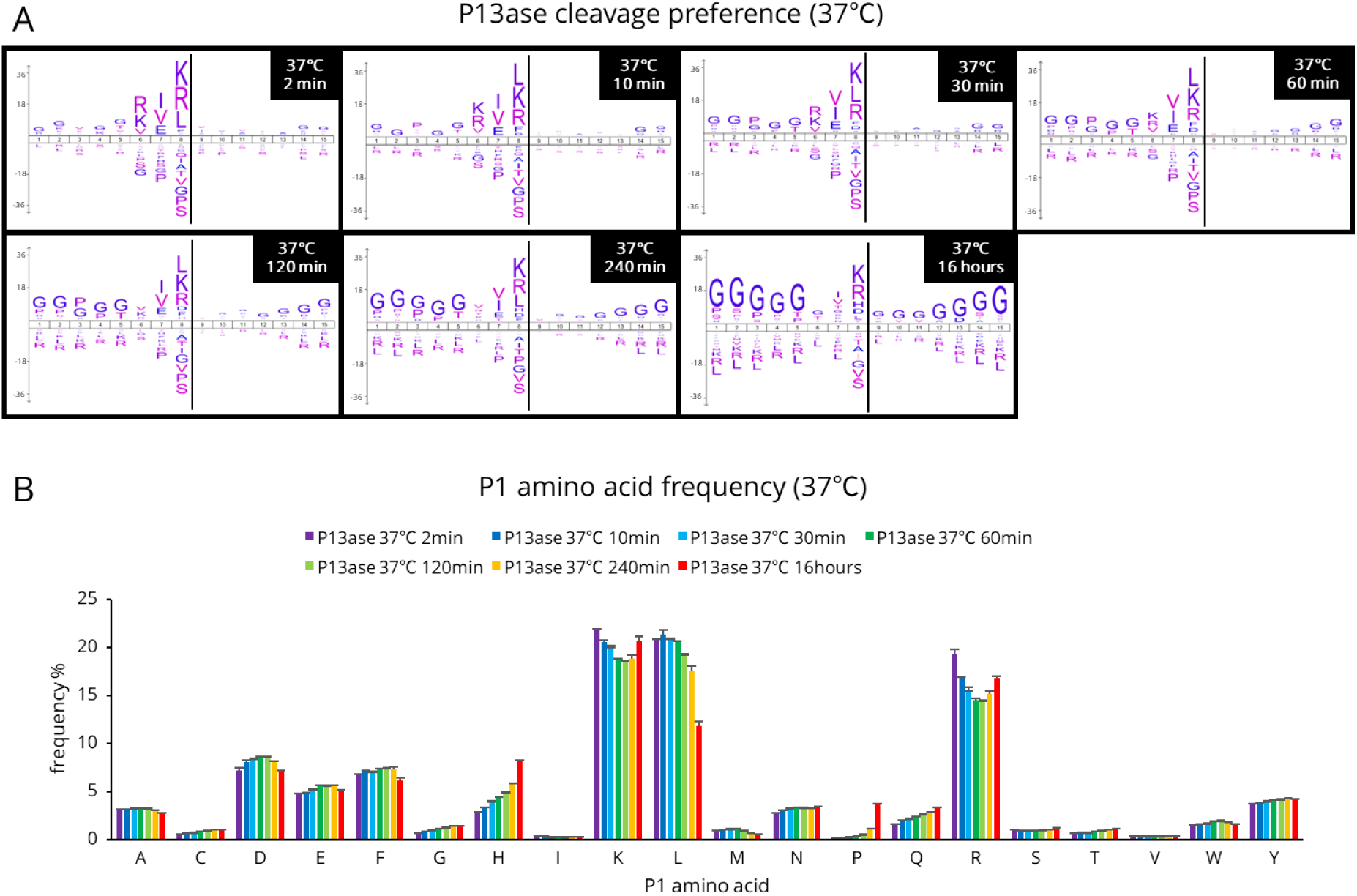
Cleavage specificity of P13ase after digestion of HeLa cell extract for various times at 37°C. (A) Cleavage preferences visualized using iceLogo at 37°C after digestion by P13ase for various times. Non-specific MaxQuant search was performed to calculate cleavage preferences against the HeLa cells lysate dataset (three replicates combined). The y-axis is the percentage difference in frequency for an amino acid in the experimental set and the reference set. (B) Frequencies of P1 amino acids at cleavage sites at 37°C after digestion by P13ase for various times. The bar graphs represent the mean of three replicates, and error bars represent the standard error.

Comparing the amino acid frequencies at the P1 position, peptides containing Leu, Lys, and Arg showed high frequencies, followed by those containing Asp, Phe, and Glu (Fig. 4B). At 37°C, the frequency of peptides containing Leu decreased significantly with increasing digestion time (especially after 16 hours), while the frequencies of peptides containing His and Pro showed a tendency to increase with increasing digestion time, especially at 16 hours. Similar trends were observed at 4°C and 25°C, but the changes were more gradual at lower digestion temperatures (Supplemental Fig. S7). When the results at each temperature were compared at the digestion time giving the maximum number of identified peptides, the cleavage preferences and amino acid frequencies at the P1 position were almost the same (Fig. 5A,B). These results suggest that digestion temperature and time affect the digestion progress but do not significantly change the cleavage preferences. Next, we analyzed the effects of digestion temperature and time on the frequency of missed cleavages. For Lys, Arg, Leu, and Glu at position P1, the percentage of peptides with missed cleavages decreased with increasing digestion temperature and time (Supplemental Fig. S8). On the other hand, the frequency of missed cleavages for Asp and Phe did not decrease significantly.

**Figure 5:**
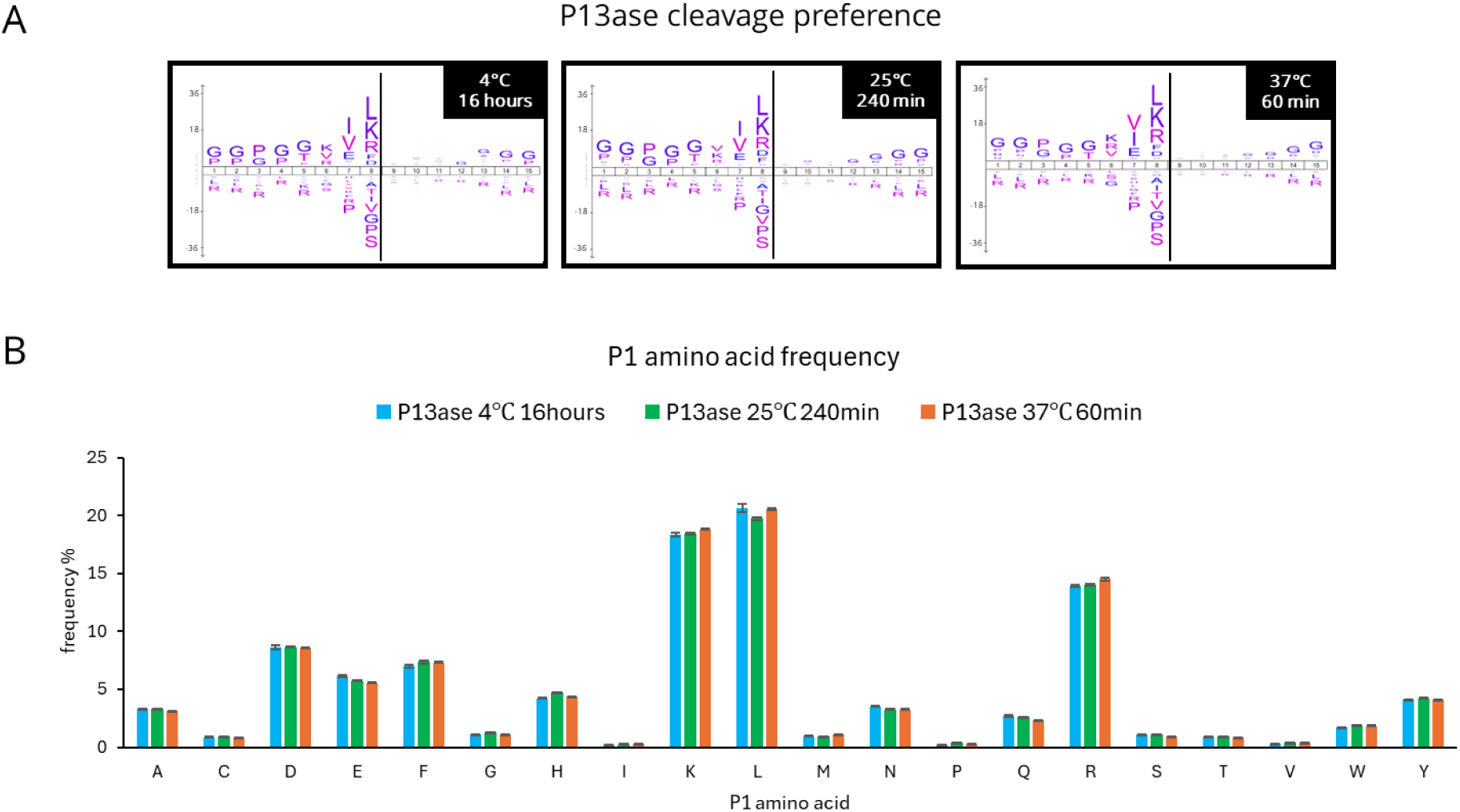
Cleavage specificity of P13ase for digestion times that maximized the number of peptides identified from HeLa cell extract at each temperature. (A) Cleavage preferences visualized using iceLogo digestion time by P13ase for digestion times that maximized the number of peptides identified at each temperature. Non-specific MaxQuant search was performed to calculate cleavage preferences against the HeLa cells lysate dataset (three replicates combined). The y-axis is the percentage difference in frequency for an amino acid in the experimental set and the reference set. (B) Frequencies of P1 amino acids at P13ase cleavage sites for digestion times that maximized the number of peptides identified at each temperature. The bar graphs represent the mean of three replicates, and error bars represent the standard error.

Next, artifacts of deamidation and succinimidation were evaluated. At all digestion temperatures and times, digestion with P13ase suppressed deamidation and succinimidation compared to trypsin (Supplemental Fig. S9). Peptides generated by self-digestion were also compared to those in the case of trypsin. The trypsin used in this study was methylated at lysine, which is believed to suppress autolysis to some extent. However, despite the protease concentration being higher than that of trypsin, the proportion of autolysis peptides for P13ase was mostly less than one-tenth of that for trypsin (Supplemental Fig. S10). However, it is currently unclear whether P13ase undergoes a change in selectivity due to self-digestion, analogous to the conversion of trypsin to show chymotrypsin-like activity.

Based on the above results, the digestion conditions for P13ase in bottom-up proteomics were selected as pH 3.5, 37 °C, and 60 minutes.

### Characterization of peptides and proteins identified by P13ase digestion

Peptides and proteins generated by P13ase digestion under conditions optimized for bottom-up proteomics (pH 3.5, 37°C, 60 min digestion) were characterized. We found that few peptides were commonly identified by P13ase and trypsin. This may be due to differences in the cleavage preferences of the proteases (Supplemental Fig. S11A). The physico-chemical properties including *m/z*, pI, molecular weight and gravy score for peptides identified by P13ase and trypsin were compared, and no significant difference was found (Supplemental Fig. S11B). The median relative standard deviation of the quantitative reproducibility of the identified unique peptides was 10.5% for P13ase, which is sufficient for general bottom-up proteomics (Supplemental Fig. S11C). On the other hand, most of the proteins identified by P13ase digestion overlapped with those identified by trypsin digestion (Supplemental Fig. S12A). There was no significant difference in the distribution of pI or molecular weight of the proteins identified by trypsin digestion and P13ase digestion (Supplemental Fig. S12B). The quantitative reproducibility of the identified proteins was also evaluated. The median relative standard deviation of quantified proteins with P13ase digestion was 9.6%, which is sufficient for general bottom-up proteomics (Supplemental Fig. S12C). Although the peptides identified by P13ase digestion and trypsin digestion shared few common peptides due to the different cleavage preferences, there was no obvious bias in the properties of the identified peptides and proteins, and quantitative reproducibility was sufficient. These results confirm the suitability of P13ase for application to bottom-up proteomics.

The number of peptides identified per protein was higher in the case of P13ase digestion than trypsin digestion (Supplementary Figure S12D). Comparison of protein sequence coverage also showed that trypsin gave higher values for proteins with relatively low sequence coverage, but P13ase gave higher values for proteins with relatively high sequence coverage (Supplementary Fig. S12E). In particular, the number of proteins with sequence coverage greater than 90% was 37 in the case of P13ase digestion compared to only 2 for trypsin digestion. As a representative example, trypsin digestion of histone H2A type 1 (Uniprot accession P0C0S8) yielded 46% coverage and failed to identify K/R-rich and K/R-poor regions (Supplementary Fig. S13). This was because the peptides produced by trypsin were either too long or too short to be identified. In contrast, P13ase digestion yielded 95% coverage. The reason for the high coverage was thought to be that P13ase does not necessarily cleave the C-terminal side of all Lys and Arg residues, allowing identification of K/R-rich regions. In addition, P13ase cleaved the C-terminal side of Leu, allowing identification of K/R-poor regions.

### Reduction of modification artifacts

As mentioned above, the rapid digestion by P13ase under acidic conditions is expected to reduce the deamidation artifact, compared with trypsin digestion. Here we analyze the results in more detail. First, the number of peptides containing deamidation modifications was less than 7% in P13ase digestion compared to trypsin digestion (Figure 6A). Of the deamidation sites identified, only 11 were common (Fig. 6B), possibly because the sites identified only with P13ase were too long when contained in tryptic peptides. For quantitative comparison, we focused on one peptide from glyceraldehyde-3-phosphate dehydrogenase as a commonly identified peptide containing the same deamidation site. In trypsin digestion, the peak area indicated that approximately 1% of the peptide was deaminated (Fig. 6C). In contrast, P13ase digestion yielded a peak area for unmodified peptide similar to that of trypsin digestion, but the deamidated peptide was below the limit of quantitation. These results suggest that P13ase, which produces fewer artifacts during digestion, is appropriate for deamidation proteomics. Modifications other than Asn deamidation were similarly examined (Fig. 6D). For both Asn and Gln, P13ase digestion reduced the frequency of deamidation, succinimidation and pyroglutamination. As for N-terminal cyclization of carbamidomethyl-cysteine, there was little difference between trypsin and P13ase. In contrast, N-terminal cyclization of Glu was less frequent with P13ase, and N-terminal cyclization of Asp was almost absent. These results suggest that P13ase digestion inhibits not only post-cyclization hydrolysis of Asn, but also cyclization and post-cyclization hydrolysis of other amino acids under acidic conditions. Thus, P13ase is advantageous in that it can be conducted under acidic conditions.

**Figure 6:**
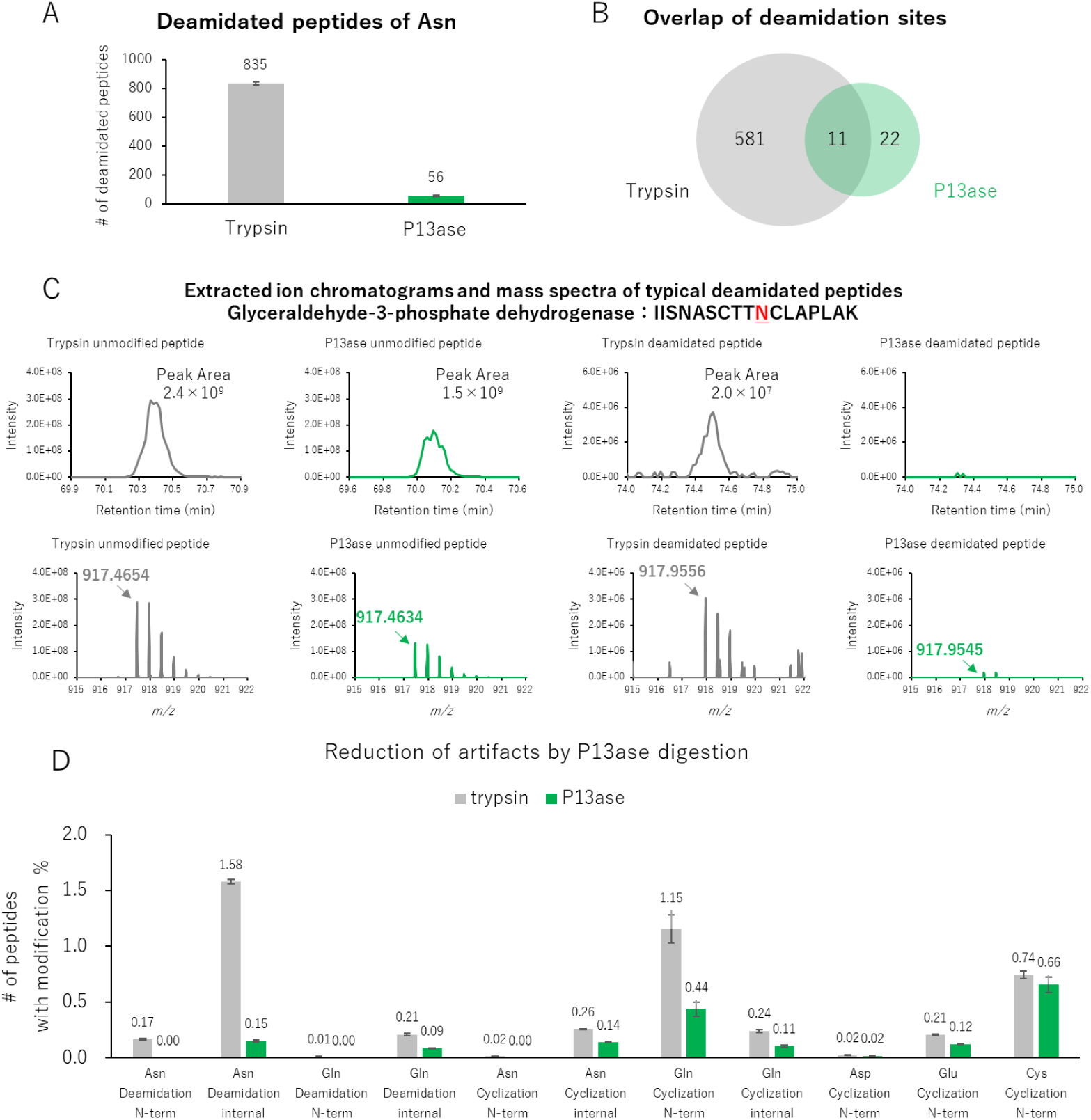
Reduction of artifacts of deamidation and cyclization by P13ase digestion of HeLa cell extract. (A) The number of identified Asn-deamidated peptides generated by trypsin (37°C, 16 hours) or P13ase (37°C, 60 min) digestion. The bar graphs represent the mean of three replicates, and error bars represent the standard error. (B) Overlap of identified deamidation sites generated by trypsin and P13ase digestion. (C) Extracted ion chromatograms (XIC) and mass spectra of typical deamidated and unmodified peptides (glyceraldehyde-3-phosphate dehydrogenase: IISNASCTTNCLAPLAK). XIC and mass spectra were analyzed by FreeStyle (Thermo Fisher Scientific). (D) Ratios of the number of identified peptides deamidated or cyclized at each amino acid after digestion with trypsin (37°C, 16 hours) or P13ase (37°C, 60 minutes). “N-term” in the bar graph means N-terminal modification of the identified peptide, and “internal” means internal modification of the identified peptide. Deamidation (NQ), cyclization (NQ) and peptide N-terminal cyclization (CDE) were added as variable modifications and searched individually.

Acidic conditions also inhibit the rearrangement of disulfide bonds (i.e., disulfide bond scrambling), which may enable the analysis of proteins without the need for alkylation by IAA or other alkylating agents. Since it has been reported that alkylating agents react with amino acids other than Cys, avoiding the use of alkylating agents may enable the capture of the true state of proteins. Additionally, while P13ase has not yet surpassed trypsin in terms of identification efficiency, there is still potential for improving identification and quantification performance through the application of data-independent acquisition and modifications to the open search method. These points will be investigated in future studies.

## CONCLUSION

In this study, we investigated the potential utility of P13ase as an alternative or complementary protease to trypsin in bottom-up proteomics. We evaluated the effects of factors such as pH, temperature, and time of digestion on the cleavage efficiency and selectivity of P13ase, and identified pH 3.5, 37 °C, and 60 minutes as the optimal conditions for bottom-up proteomics. We also found that P13ase has co-occurrence motifs not only at P1 but also at other positions. Furthermore, we demonstrated that P13ase digestion can be achieved rapidly under low pH conditions, thereby minimizing deamidation and other artifacts. These results not only extend the applicability of P13ase in standard bottom-up proteomics workflows, but also suggest its suitability for applications requiring artifact-free proteomics in order to target only intrinsic modifications.

## Acknowledgment

This work was supported by JST SPRING (JPMJSP2123 to AT), JST ACT-X (JPMJAX2324 to KO), JSPS KAKENHI (23K18185, 23H04924 to YI, 24KJ1440 to AT, and 25K18607 to KO), FY 2024 Researcher Exchange Program between JSPS and ETH to KO, Nakatani Foundation to KO and the Research Foundation for Pharmaceutical Sciences to KO.

## Author contributions

Y. I. Conceptualization. R.T., A.T., K.O., and Y. I. designed experiments. R.T. performed experiments. R.T. and A.T. analyzed data. Y.I. supervised the project and provided resources for the study. R.T. and Y. I. wrote the paper. Approval of the final version of the paper was carried out by all authors.

## Supporting Information

**Supplementary Figure S1.**
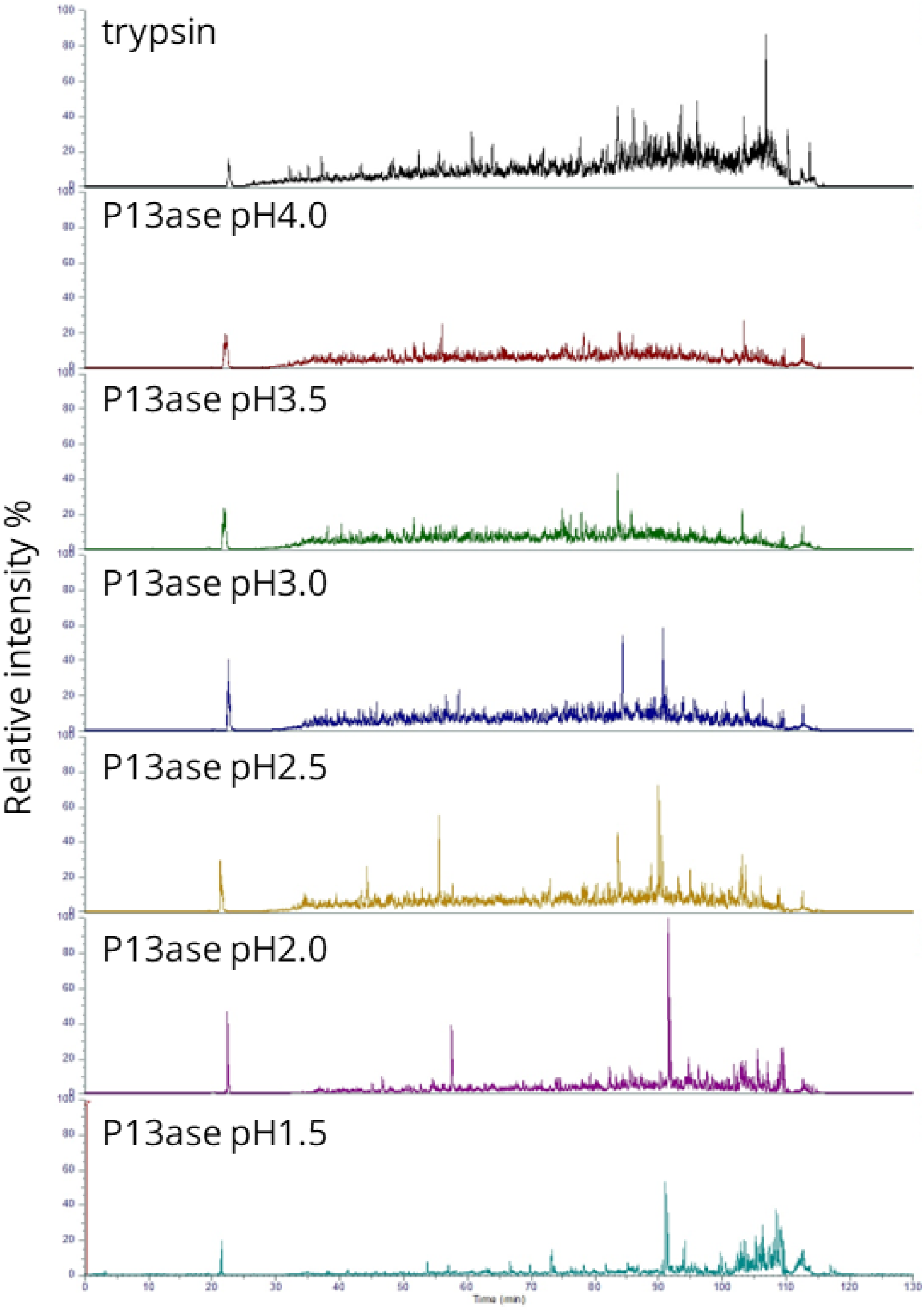
Total ion chromatograms after P13ase digestion of HeLa cell extract at various pH values. Total ion chromatograms of digests obtained with P13ase at various pH values (pH 1.5 - 4.0) or with trypsin (pH 9.2). Chromatograms were analyzed with FreeStyle (Thermo Fisher Scientific).

**Supplementary Figure S2.**
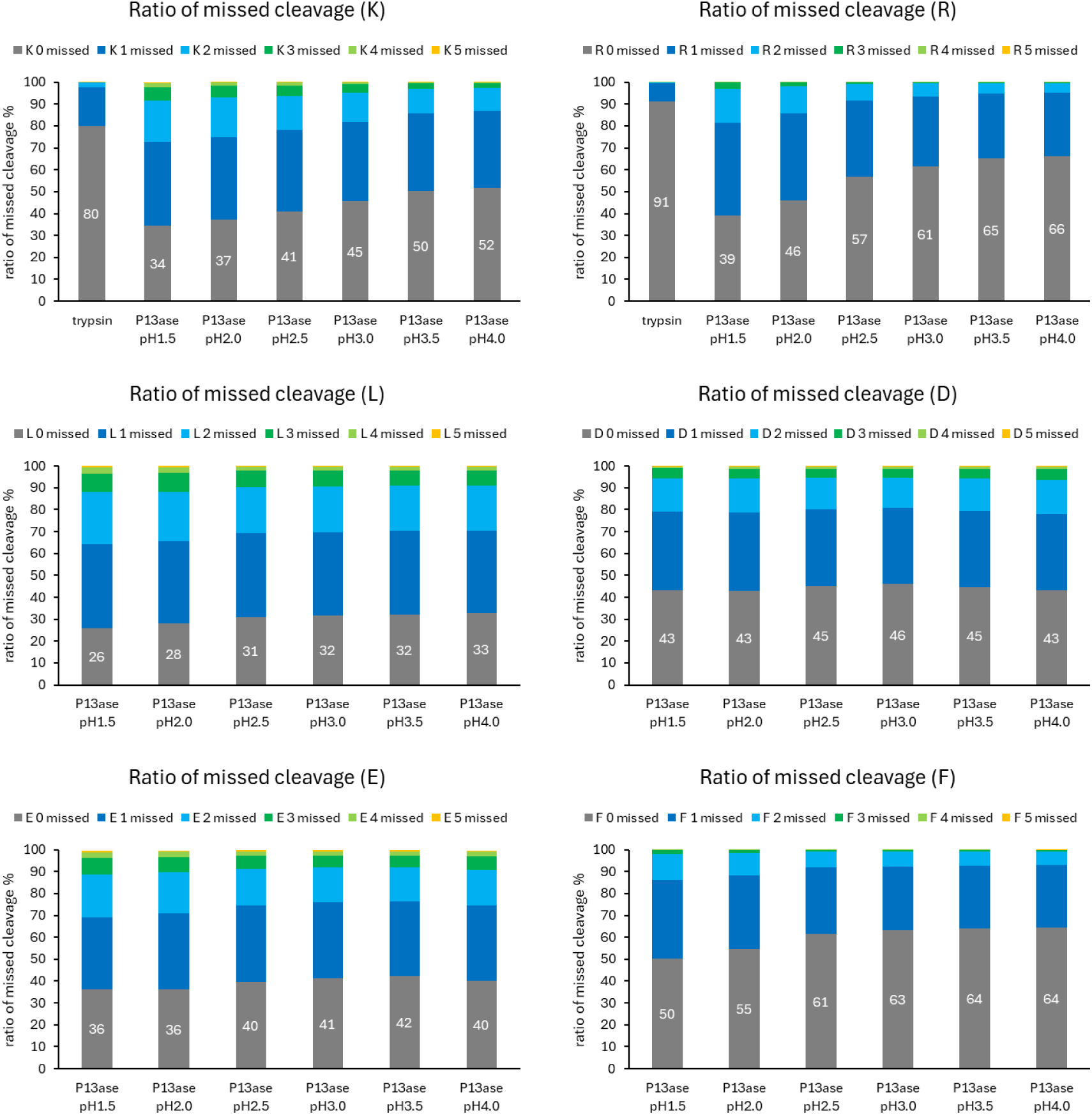
Frequencies of missed cleavage by P13ase at various pH values. The missed cleavage frequencies of Lys, Arg, Leu, Phe, Asp and Glu residues by P13ase at various pH values (1.5 - 4.0). Frequencies were averaged from three technical replicates.

**Supplementary Figure S3.**
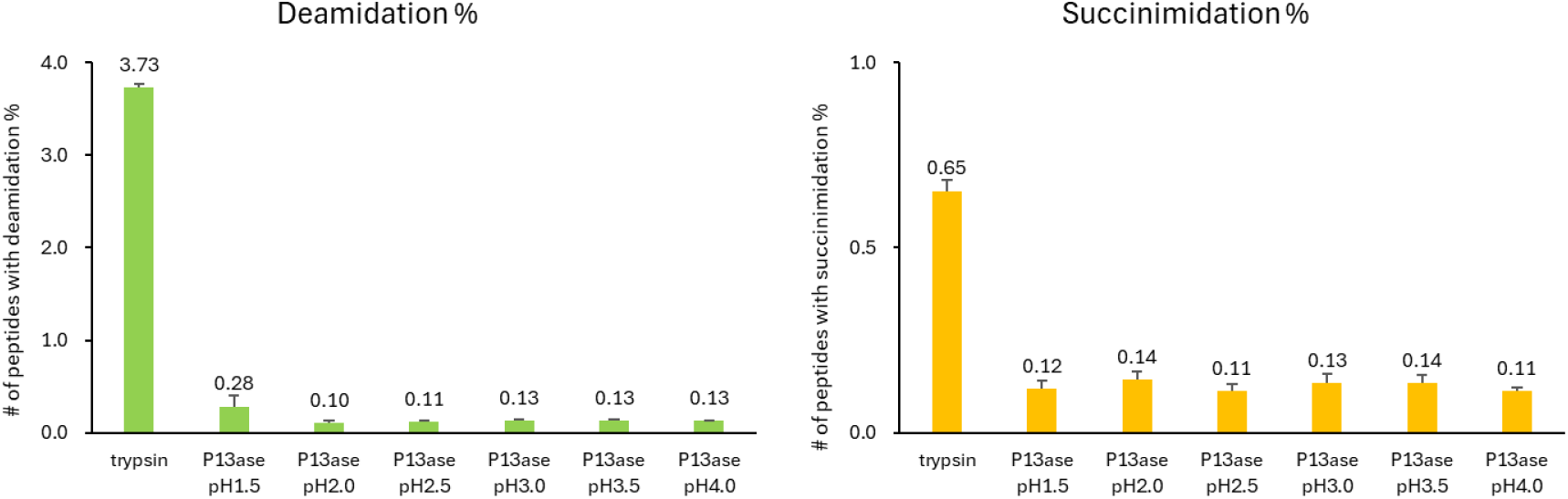
Effect of digestion pH on artifactual deamidation and succinimidation. Comparison of the proportion of peptides containing deamidation or succinimidation after digestion with P13ase or trypsin at various pH values. The bar graphs represent the mean of three technical replicates and error bars represent the standard error.

**Supplementary Figure S4.**
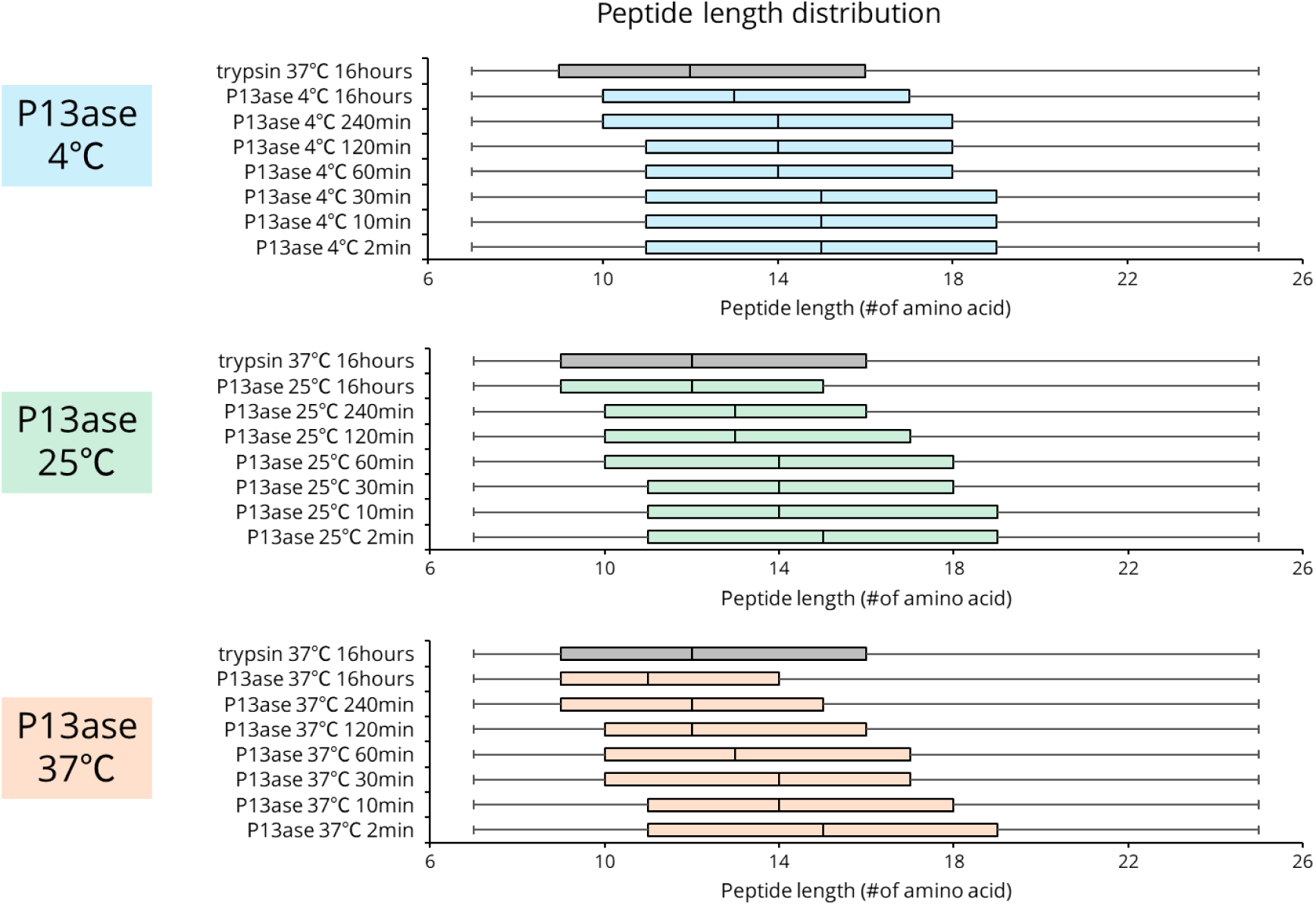
Peptide lengths after P13ase digestion of HeLa cell extract for various times and at various temperatures. The length distribution of identified peptides after P13ase digestion for various times and at various temperatures.

**Supplementary Figure S5.**
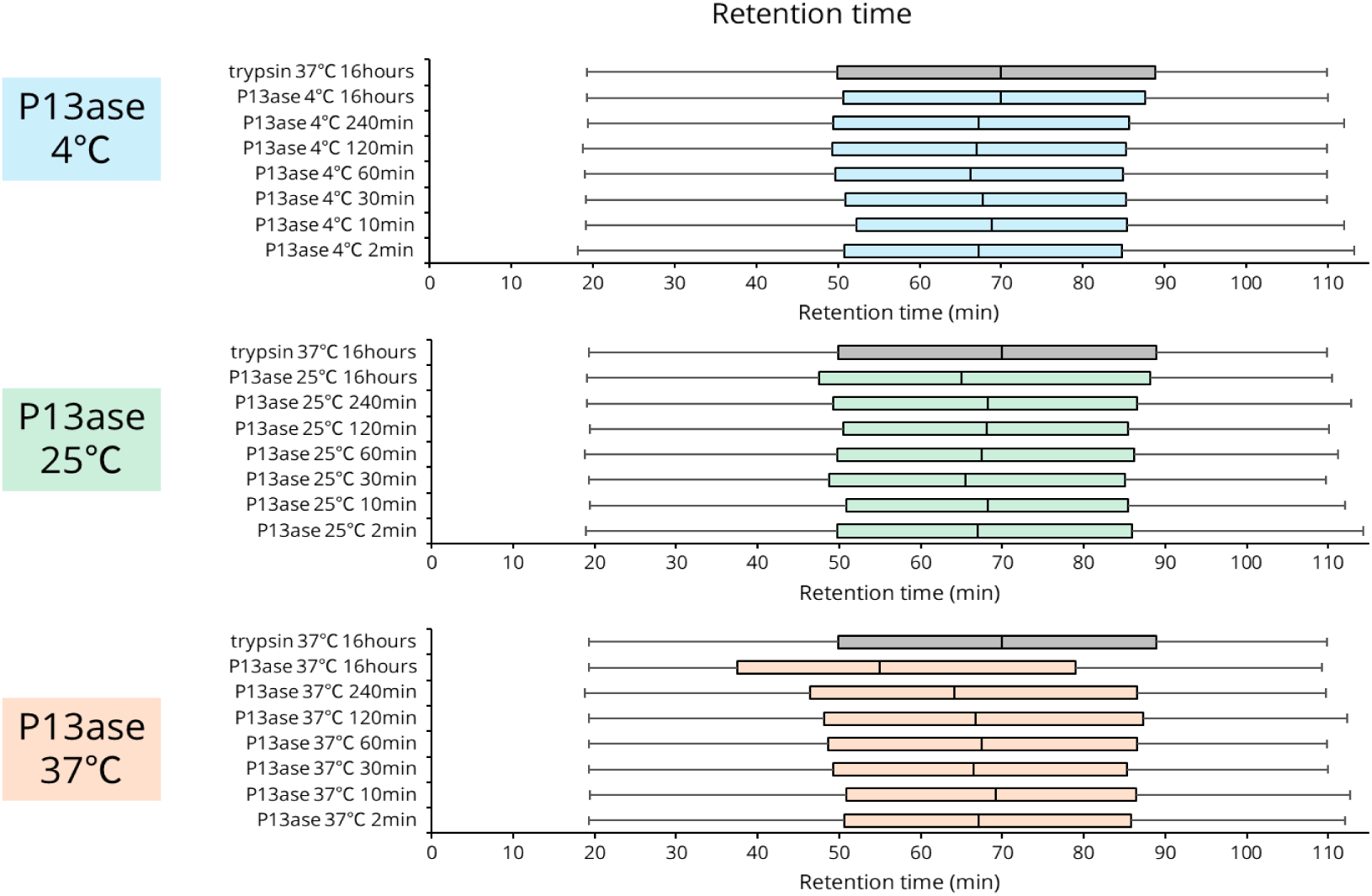
Retention time distribution after P13ase digestion of HeLa cell extract for various times and at various temperatures. The distribution of retention times of identified peptides after P13ase digestion for various times and at various temperatures.

**Supplementary Figure S6.**
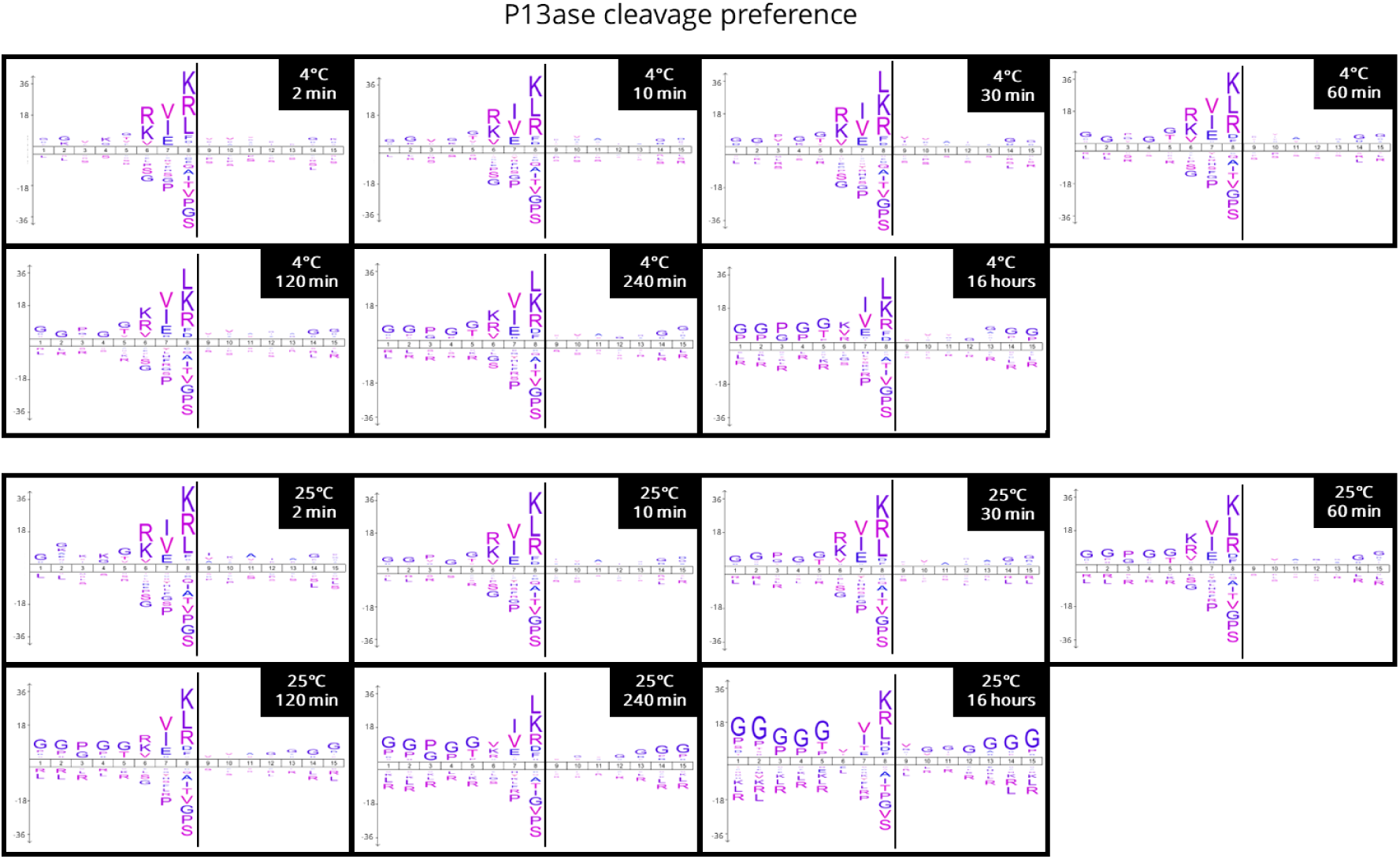
Cleavage preferences of P13ase at various digestion times and temperatures. Cleavage preferences were visualized with iceLogo for various digestion times and temperatures. The y-axis is the percentage difference in frequency for an amino acid in the experimental set and the reference set.

**Supplementary Figure S7.**
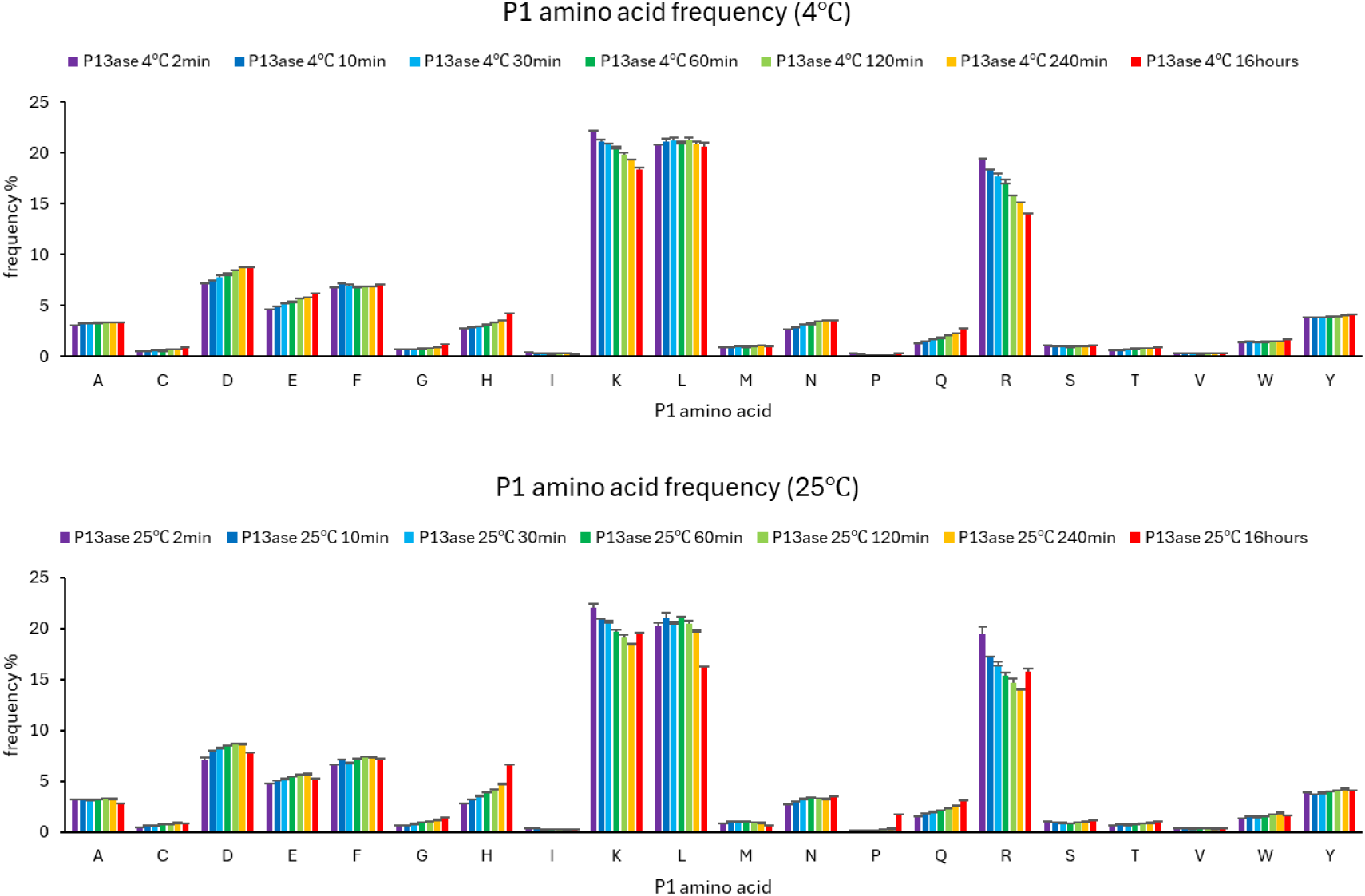
Cleavage specificity of P13ase at various digestion temperatures and times. Frequencies of P1 amino acids at cleavage sites after digestion with P13ase at various temperatures for various times. The bar graphs represent the mean of three technical replicates and error bars represent the standard error.

**Supplementary Figure S8.**
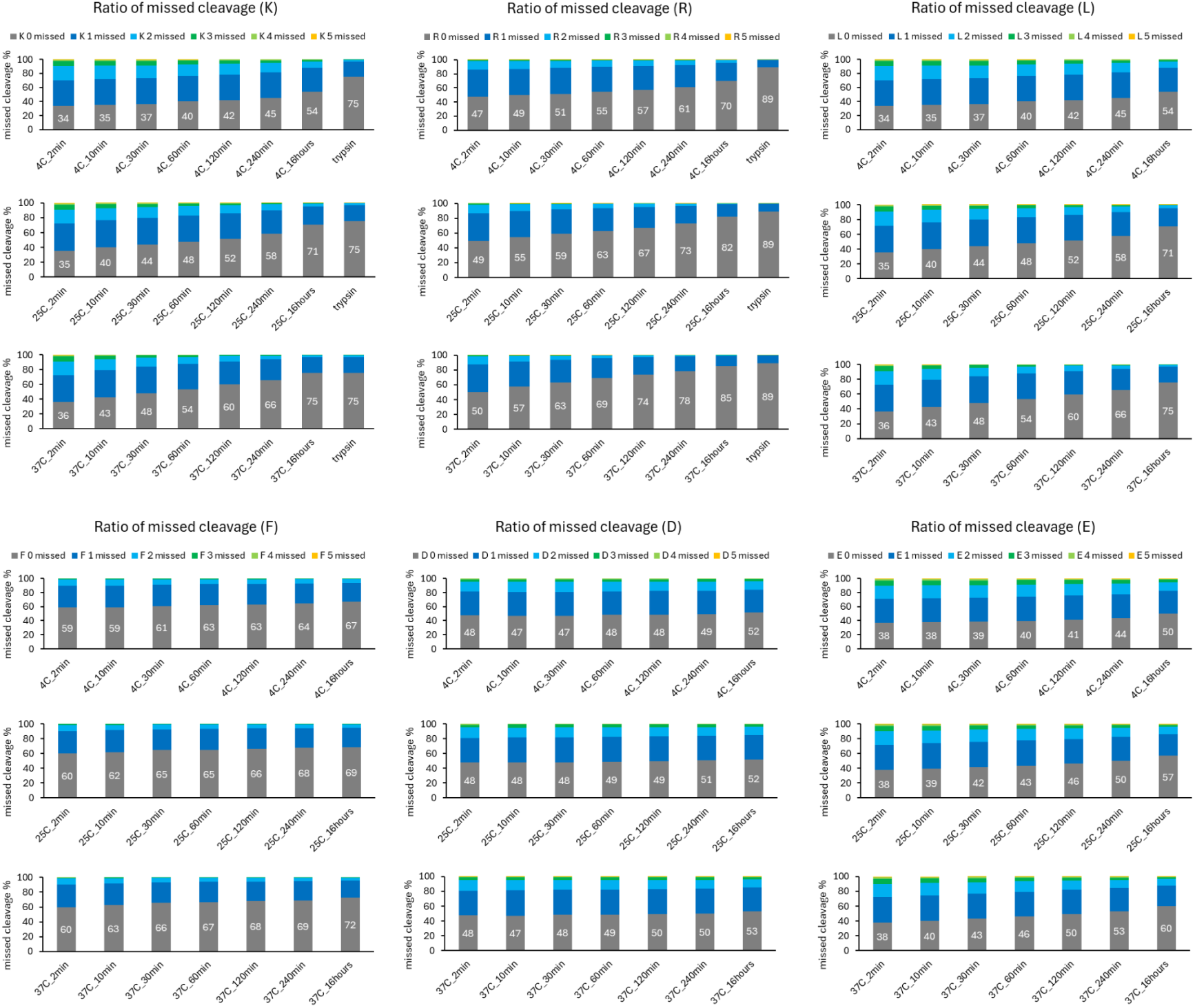
Missed cleavage frequencies of P13ase at various digestion temperatures and times. The missed cleavage frequencies of Lys, Arg, Leu, Phe, Asp and Glu residues by P13ase at various digestion temperatures and times. Frequencies were averaged from three technical replicates.

**Supplementary Figure S9.**
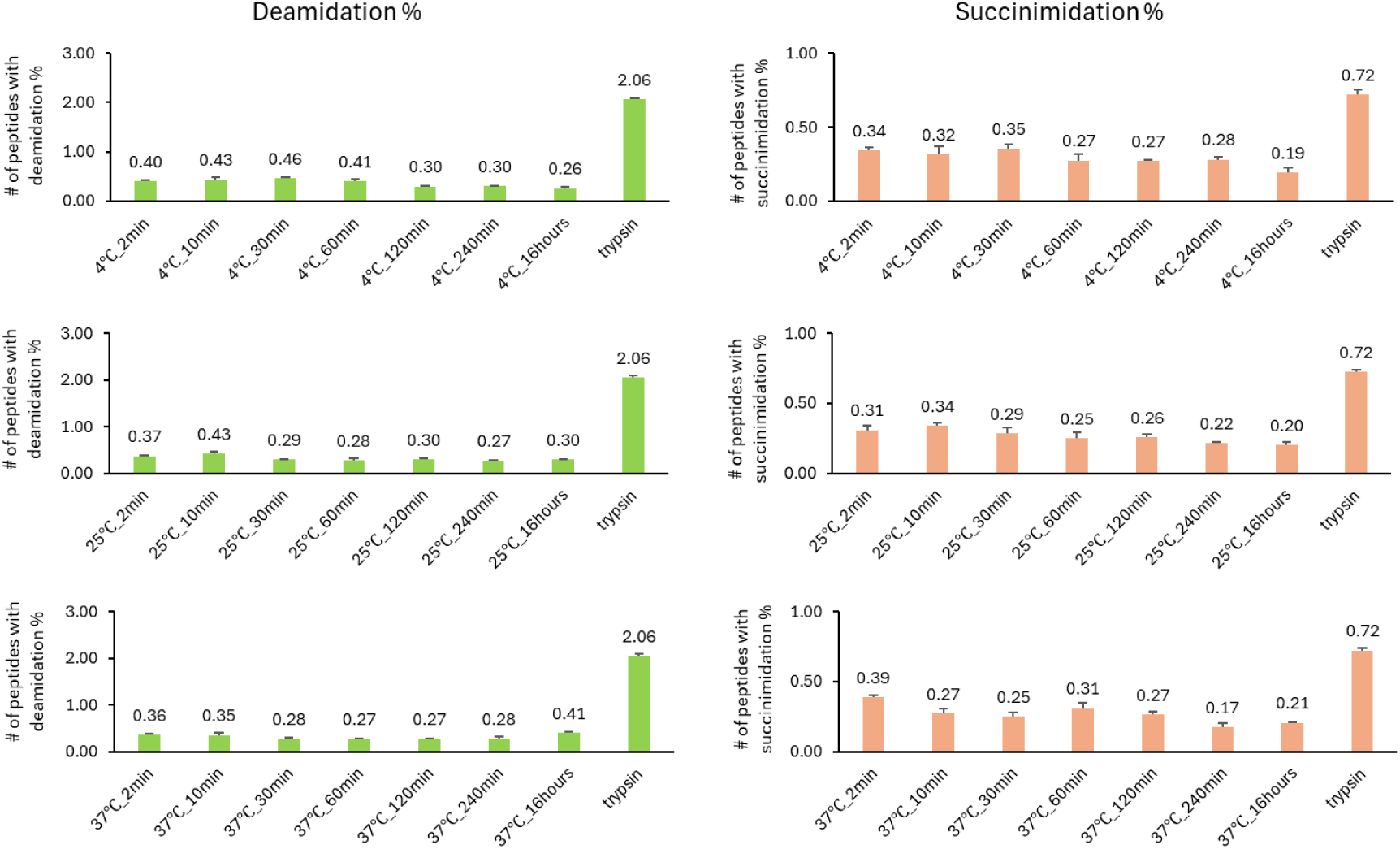
Effect of digestion temperature and time on artifactual deamidation and succinimidation. Comparison of the proportion of peptides containing deamidation or succinimidation after digestion with P13ase or trypsin at various temperatures and for various times. The bar graphs represent the mean of three technical replicates and error bars represent the standard error.

**Supplementary Figure S10.**
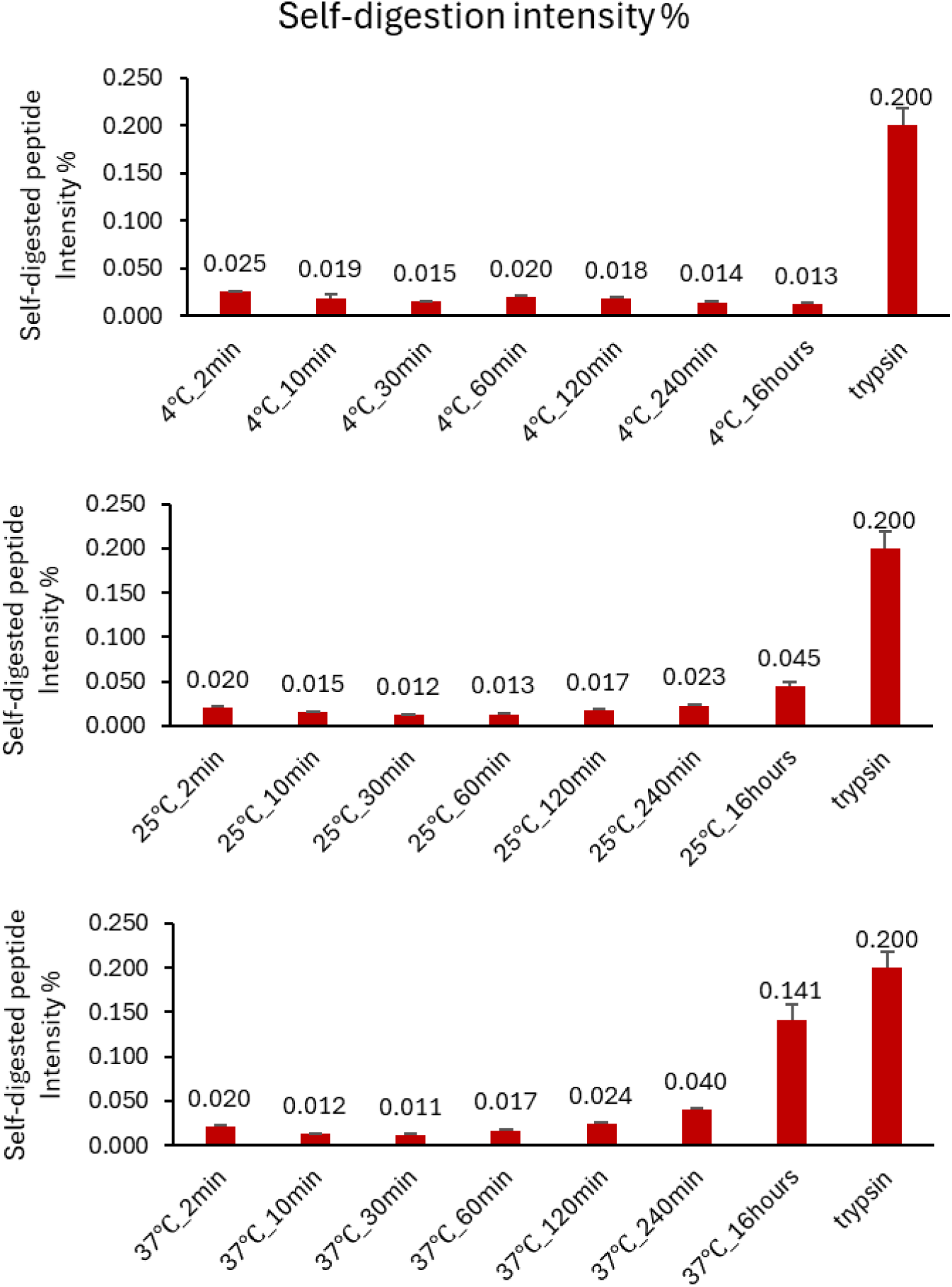
Self-digested peptides generated by P13ase or trypsin. The ratios of summed intensity of self-digested identified peptides produced by P13ase or trypsin at various digestion temperatures and for various times. The bar graphs represent the mean of three technical replicates and error bars represent the standard error.

**Supplementary Figure S11.**
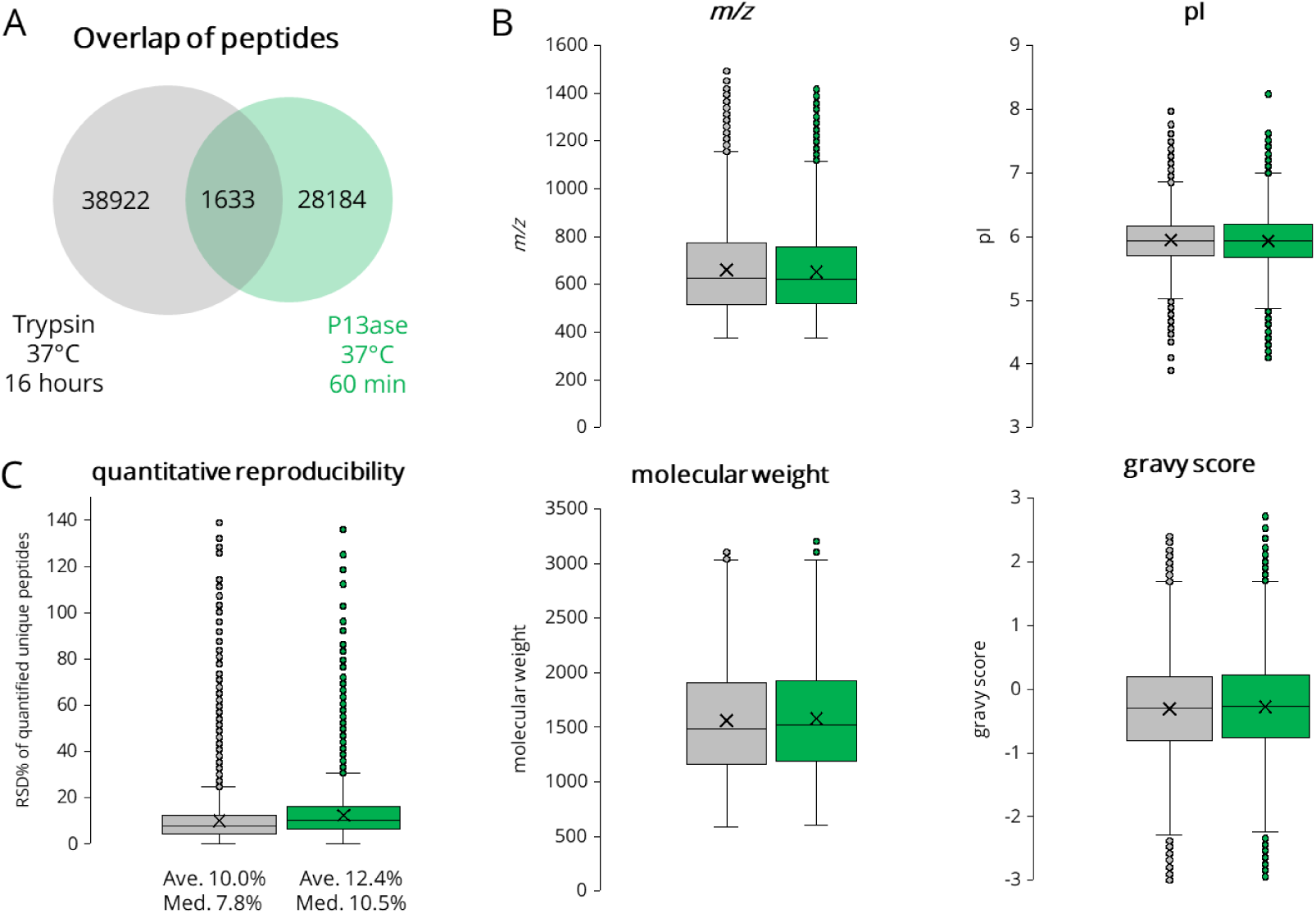
Characteristics of peptides identified by P13ase digestion of HeLa cell extract. (A) Overlap of peptides identified by P13ase (37°C, 60 min digestion) and trypsin (37°C, 16 hours digestion). (B) Comparison of physicochemical properties (*m/z*, pI, molecular weight, gravy score) of peptides identified by P13ase and trypsin. (C) Quantitative reproducibility of identified peptides.

**Supplementary Figure S12.**
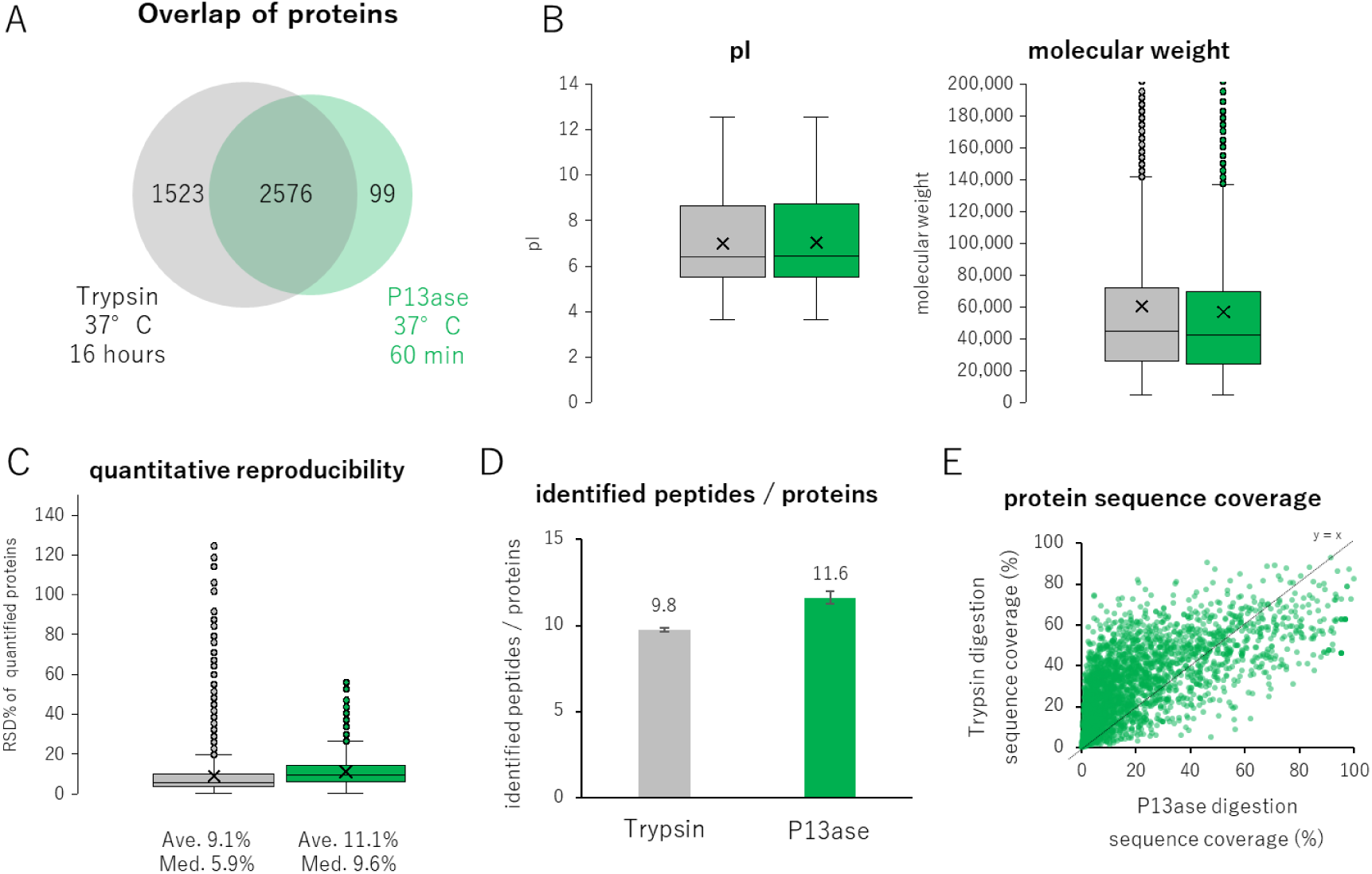
Characteristics of proteins identified by P13ase digestion of HeLa cell extract. (A) Overlap of proteins identified by digestion with P13ase (37°C, 60 min) and trypsin (37°C, 16 hours). (B) Distributions of pI and molecular weights of proteins identified by P13ase and trypsin digestion. (C) Quantitative reproducibility of proteins identified by P13ase and trypsin digestion. (D) The number of peptides identified per protein by P13ase and trypsin digestion. The bar graphs represent the mean of three replicates, and error bars represent the standard error. (E) Scatterplot of sequence coverage of proteins commonly identified by P13ase and trypsin digestion. The sequence coverages were calculated with Protein Coverage Summarizer.

**Supplementary Figure S13.**
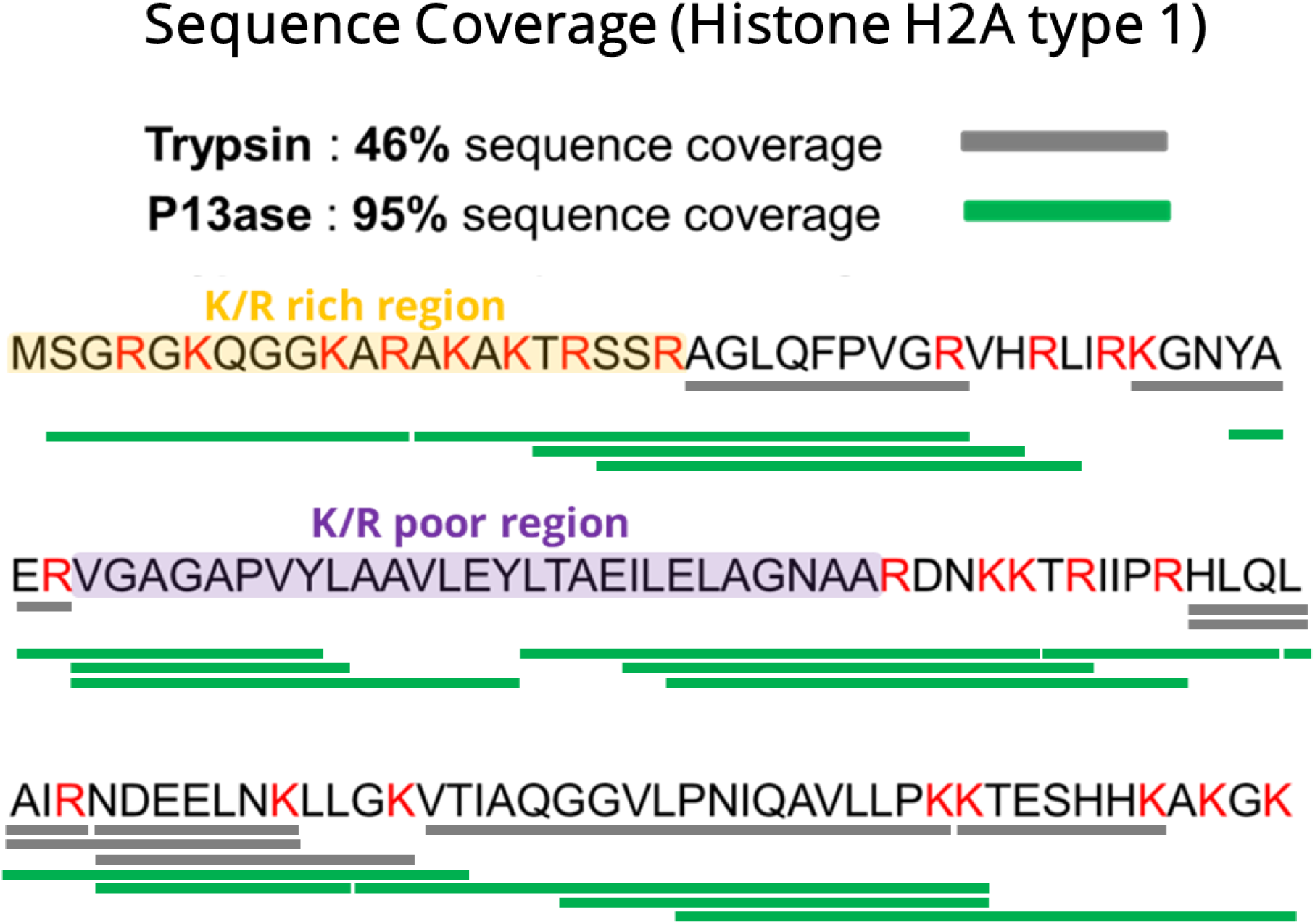
Sequence coverage of histone H2A type 1 by P13ase or trypsin digestion. Sequence coverage of histone H2A type 1 by P13ase (37°C, 60 min digestion) or trypsin (37°C, 16 hours digestion). The sequence coverages were calculated with Protein Coverage Summarizer.

**Supplementary Table S1.**
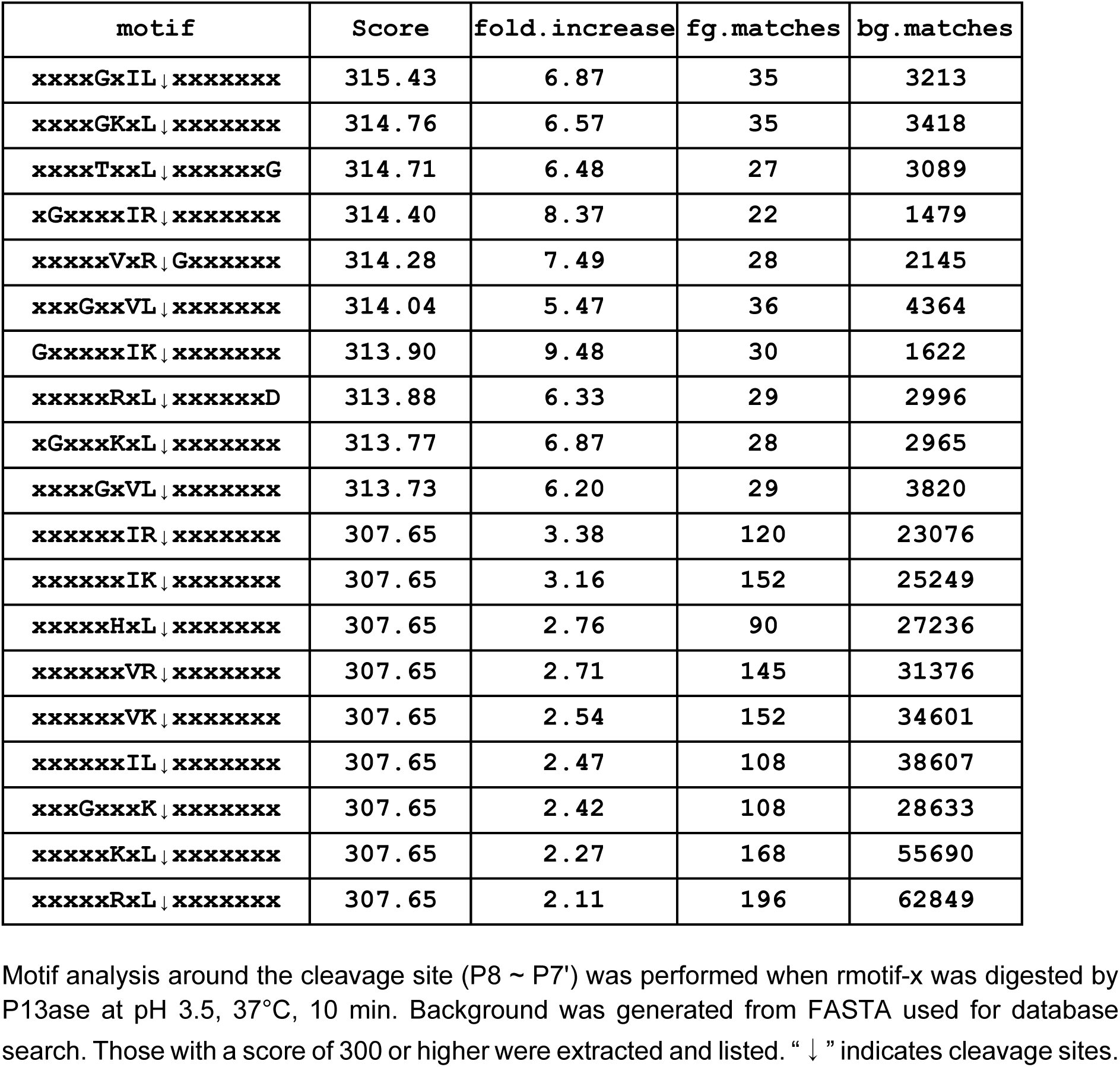
Motif analysis of the cleavage sites of P13ase (pH 3.5)

**Supplementary Table S2.**
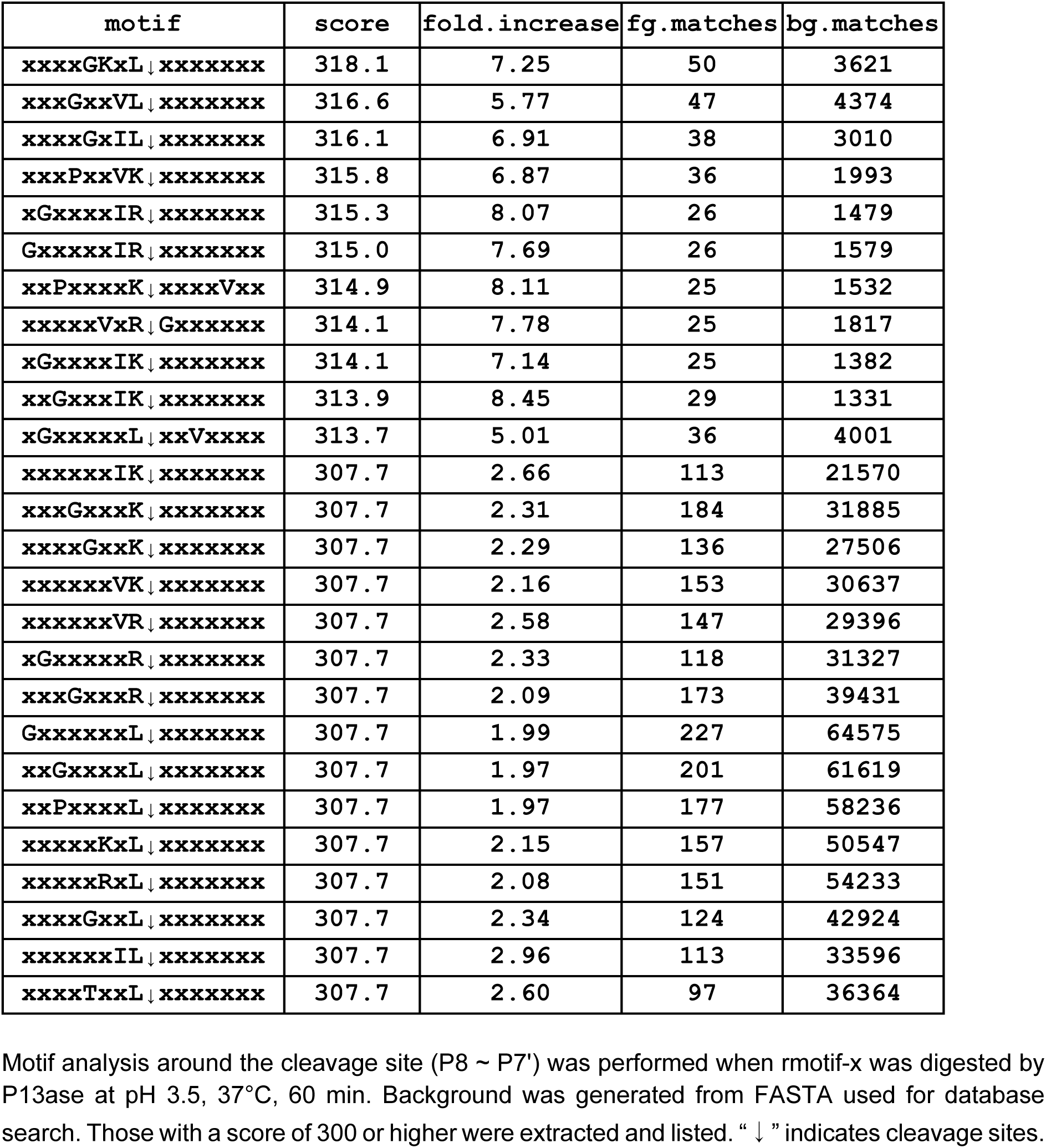
Motif analysis of the cleavage sites of P13ase (37°C, 60 minutes)

**Supplementary Table S3.**
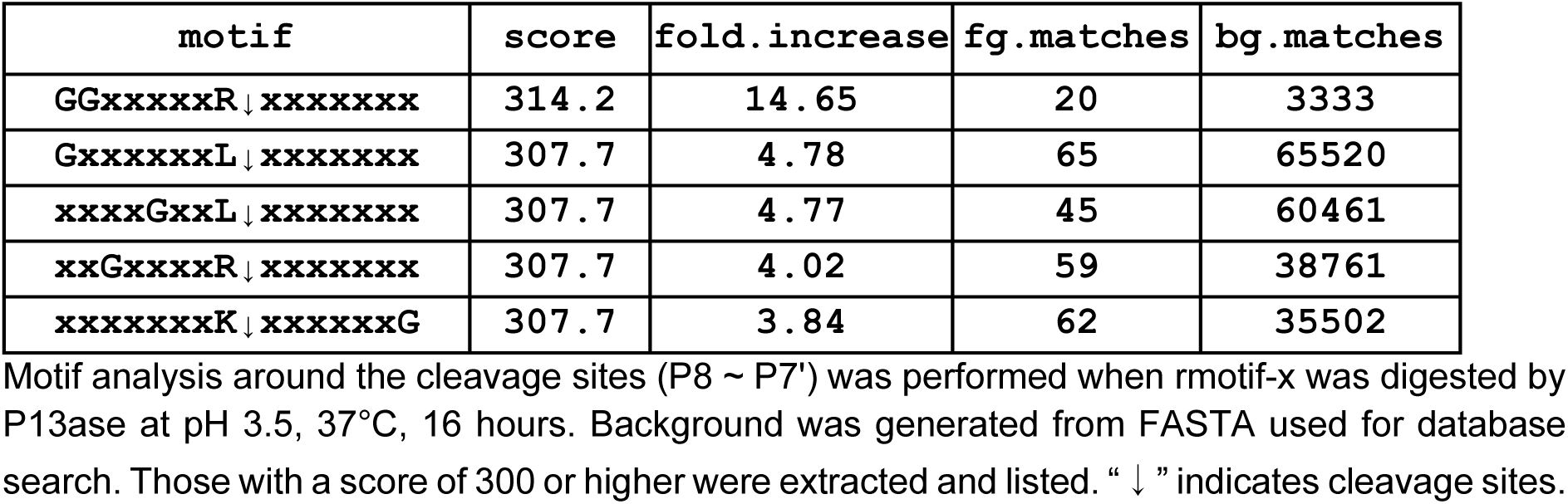
Motif analysis of the cleavage site of P13ase (37°C, 16 hours)

## REFERENCES

1. Dick, L. W., Jr, Mahon, D., Qiu, D., and Cheng, K.-C. (2009) Peptide mapping of therapeutic monoclonal antibodies: improvements for increased speed and fewer artifacts. J. Chromatogr. B Analyt. Technol. Biomed. Life Sci. 877, 230–236

2. Aebersold, R., and Mann, M. (2016) Mass-spectrometric exploration of proteome structure and function. Nature 537, 347–355

3. Müller, J. B., Geyer, P. E., Colaço, A. R., Treit, P. V., Strauss, M. T., Oroshi, M., Doll, S., Virreira Winter, S., Bader, J. M., Köhler, N., Theis, F., Santos, A., and Mann, M. (2020) The proteome landscape of the kingdoms of life. Nature 582, 592–596

4. Tsiatsiani, L., and Heck, A. J. R. (2015) Proteomics beyond trypsin. FEBS J. 282, 2612–2626

5. Sinitcyn, P., Richards, A. L., Weatheritt, R. J., Brademan, D. R., Marx, H., Shishkova, E., Meyer, J. G., Hebert, A. S., Westphall, M. S., Blencowe, B. J., Cox, J., and Coon, J. J. (2023) Global detection of human variants and isoforms by deep proteome sequencing. Nat. Biotechnol.,

6. Giansanti, P., Tsiatsiani, L., Low, T. Y., and Heck, A. J. R. (2016) Six alternative proteases for mass spectrometry-based proteomics beyond trypsin. Nat. Protoc. 11, 993–1006

7. Samodova, D., Hosfield, C. M., Cramer, C. N., Giuli, M. V., Cappellini, E., Franciosa, G., Rosenblatt, M. M., Kelstrup, C. D., and Olsen, J. V. (2020) ProAlanase is an Effective Alternative to Trypsin for Proteomics Applications and Disulfide Bond Mapping. Mol. Cell. Proteomics 19, 2139–2157

8. Muriithi, B., Ippoliti, S., Finny, A., Addepalli, B., and Lauber, M. (2024) Clean and complete protein digestion with an autolysis resistant trypsin for peptide mapping. J. Proteome Res. 23, 5221–5228

9. Longarini, E. J., and Matić, I. (2024) Preserving ester-linked modifications reveals glutamate and aspartate mono-ADP-ribosylation by PARP1 and its reversal by PARG. Nat. Commun. 15, 4239

10. Jiang, X., Yeung, D., Liu, Y., Spicer, V., Afshari, H., Lao, Y., Lin, F., Krokhin, O., and Zahedi, R. P. (2024) Accelerating Proteomics Using Broad Specificity Proteases. J. Proteome Res.,

11. Tomioka, R., Tomioka, A., Ogata, K., Chan, H.-J., Chen, L.-Y., Guzman, U. H., Xuan, Y., Olsen, J. V., Chen, Y.-J., and Ishihama, Y. (2024) Extending the coverage of Lys-C/trypsin-based bottom-up proteomics by cysteine S-aminoethylation. J. Am. Soc. Mass Spectrom. 35, 386–396

12. Iwasaki, M., Masuda, T., Tomita, M., and Ishihama, Y. (2009) Chemical cleavage-assisted tryptic digestion for membrane proteome analysis. J. Proteome Res. 8, 3169–3175

13. Ofori, S., Desai, H. S., Shikwana, F., Boatner, L. M., Dominguez, E. R., Iii Castellón, J. O., and Backus, K. M. (2024) Generating cysteine-trypsin cleavage sites with 2- chloroacetamidine capping. Chem. Commun. (Camb.) 60, 8856–8859

14. Ying, Y., and Li, H. (2022) Recent progress in the analysis of protein deamidation using mass spectrometry. Methods 200, 42–57

15. Scotchler, J. W., and Robinson, A. B. (1974) Deamidation of glutaminyl residues: dependence on pH, temperature, and ionic strength. Anal. Biochem. 59, 319–322

16. Cao, M., Xu, W., Niu, B., Kabundi, I., Luo, H., Prophet, M., Chen, W., Liu, D., Saveliev, S. V., Urh, M., and Wang, J. (2019) An automated and qualified platform method for site-specific succinimide and deamidation quantitation using low-pH peptide mapping. J. Pharm. Sci. 108, 3540–3549

17. 17. Yannone, S. M., Tuteja, V., Goleva, O., Leung, D. Y. M., Stotland, A., Keoseyan, A. J., Hendricks, N. G., Parker, S., Van Eyk, J. E., and Kreimer, S. (2025) Toward real-time proteomics: Blood to biomarker quantitation in under one hour. Anal. Chem.,

18. McCabe, M. C., Gejji, V., Barnebey, A., Siuzdak, G., Hoang, L. T., Pham, T., Larson, K. Y., Saviola, A. J., Yannone, S. M., and Hansen, K. C. (2023) From volcanoes to the bench: Advantages of novel hyperthermoacidic archaeal proteases for proteomics workflows. J. Proteomics 289, 104992

19. Tomioka, R., Ogata, K., and Ishihama, Y. (2024) Quantitation of host cell proteins by capillary LC/IMS/MS/MS in combination with rapid digestion on immobilized trypsin column under native conditions. Mass Spectrom. (Tokyo) 13, A0152

20. Masuda, T., Saito, N., Tomita, M., and Ishihama, Y. (2009) Unbiased quantitation of Escherichia coli membrane proteome using phase transfer surfactants. Mol. Cell. Proteomics 8, 2770–2777

21. Ota, S., Miyazaki, S., Matsuoka, H., Morisato, K., Shintani, Y., and Nakanishi, K. (2007) High-throughput protein digestion by trypsin-immobilized monolithic silica with pipette-tip formula. J. Biochem. Biophys. Methods 70, 57–62

22. Ichishima, E. (1970) in *Methods in Enzymology*, Methods in enzymology. (Elsevier), pp 397– 406.

23. Tanaka, N., Takeuchi, M., and Ichishima, E. (1977) Purification of an acid proteinase from Aspergillus saitoi and determination of peptide bond specificity. Biochim. Biophys. Acta 485, 406–416

24. Mullahoo, J., Zhang, T., Clauser, K., Carr, S. A., Jaffe, J. D., and Papanastasiou, M. (2020) Dual protease type XIII/pepsin digestion offers superior resolution and overlap for the analysis of histone tails by HX-MS. Methods 184, 135–140

25. Mao, Y., Zhang, L., Kleinberg, A., Xia, Q., Daly, T. J., and Li, N. (2019) Fast protein sequencing of monoclonal antibody by real-time digestion on emitter during nanoelectrospray. MAbs 11, 767–778

26. Zhang, H.-M., Kazazic, S., Schaub, T. M., Tipton, J. D., Emmett, M. R., and Marshall, A. G. (2008) Enhanced digestion efficiency, peptide ionization efficiency, and sequence resolution for protein hydrogen/deuterium exchange monitored by Fourier transform ion cyclotron resonance mass spectrometry. Anal. Chem. 80, 9034–9041

27. Cravello, L., Lascoux, D., and Forest, E. (2003) Use of different proteases working in acidic conditions to improve sequence coverage and resolution in hydrogen/deuterium exchange of large proteins. Rapid Commun. Mass Spectrom. 17, 2387–2393

28. Rey, M., Man, P., Clémençon, B., Trézéguet, V., Brandolin, G., Forest, E., and Pelosi, L. (2010) Conformational dynamics of the bovine mitochondrial ADP/ATP carrier isoform 1 revealed by hydrogen/deuterium exchange coupled to mass spectrometry. J. Biol. Chem. 285, 34981–34990

29. Rappsilber, J., Ishihama, Y., and Mann, M. (2003) Stop and go extraction tips for matrix-assisted laser desorption/ionization, nanoelectrospray, and LC/MS sample pretreatment in proteomics. Anal. Chem. 75, 663–670

30. Rappsilber, J., Mann, M., and Ishihama, Y. (2007) Protocol for micro-purification, enrichment, pre-fractionation and storage of peptides for proteomics using StageTips. Nat. Protoc. 2, 1896–1906

31. Kong, A. T., Leprevost, F. V., Avtonomov, D. M., Mellacheruvu, D., and Nesvizhskii, A. I. (2017) MSFragger: ultrafast and comprehensive peptide identification in mass spectrometry-based proteomics. Nat. Methods 14, 513–520

32. Teo, G. C., Polasky, D. A., Yu, F., and Nesvizhskii, A. I. (2021) Fast Deisotoping Algorithm and Its Implementation in the MSFragger Search Engine. J. Proteome Res. 20, 498–505

33. 33. da Veiga Leprevost, F., Haynes, S. E., Avtonomov, D. M., Chang, H.-Y., Shanmugam, A. K., Mellacheruvu, D., Kong, A. T., and Nesvizhskii, A. I. (2020) Philosopher: a versatile toolkit for shotgun proteomics data analysis. Nat. Methods 17, 869–870

34. Yu, F., Haynes, S. E., and Nesvizhskii, A. I. (2021) IonQuant Enables Accurate and Sensitive Label-Free Quantification With FDR-Controlled Match-Between-Runs. Mol. Cell. Proteomics 20, 100077

35. Cox, J., and Mann, M. (2008) MaxQuant enables high peptide identification rates, individualized p.p.b.-range mass accuracies and proteome-wide protein quantification. Nat. Biotechnol. 26, 1367–1372

36. Colaert, N., Helsens, K., Martens, L., Vandekerckhove, J., and Gevaert, K. (2009) Improved visualization of protein consensus sequences by iceLogo. Nat. Methods 6, 786–787

37. Wagih, O., Sugiyama, N., Ishihama, Y., and Beltrao, P. (2016) Uncovering phosphorylation-based specificities through functional interaction networks. Mol. Cell. Proteomics 15, 236–245

38. Okuda, S., Watanabe, Y., Moriya, Y., Kawano, S., Yamamoto, T., Matsumoto, M., Takami, T., Kobayashi, D., Araki, N., Yoshizawa, A. C., Tabata, T., Sugiyama, N., Goto, S., and Ishihama, Y. (2017) jPOSTrepo: an international standard data repository for proteomes. Nucleic Acids Res. 45, D1107–D1111

39. Li, K., Vaudel, M., Zhang, B., Ren, Y., and Wen, B. (2019) PDV: an integrative proteomics data viewer. Bioinformatics 35, 1249–1251

